# A general model for the evolution of nuptial gift-giving

**DOI:** 10.1101/2023.01.11.523385

**Authors:** Anders P. Charmouh, Trine Bilde, Greta Bocedi, A. Bradley Duthie

## Abstract

Nuptial gift-giving occurs in several taxonomic groups including insects, snails, birds, squid, arachnids and humans. Although this trait has evolved many times independently, no general framework has been developed to predict the conditions necessary for nuptial gift-giving to evolve. We use a time-in time-out model to derive analytical results describing the requirements necessary for selection to favour nuptial gift-giving. Specifically, selection will favour nuptial gift-giving if the fitness increase caused by gift-giving exceeds the product of expected gift search time and encounter rate of the opposite sex. Selection will favour choosiness in the opposite sex if the value of a nuptial gift exceeds the inverse of the time taken to produce offspring multiplied by the rate at which mates with nuptial gifts are encountered. Selection can differ between the sexes, potentially causing sexual conflict. We further investigate these results using an individual-based model inspired by a system of nuptial gift-giving spiders, *Pisaura mirabilis*, by estimating the fitness benefit of nuptial gift-giving using experimental data from several studies. Our results provide a general framework for understanding when the evolution of nuptial gift-giving can occur and provide novel insight into the evolution of worthless nuptial gifts, occurring in multiple taxonomic groups with implications for understanding parental investment.

## Introduction

Nuptial gift-giving occurs when the choosy sex (usually the female) receives gifts from the opposite sex (usually the male) during courtship. It is a widespread phenomenon, occurring within several diverse taxonomic groups such as insects, snails, birds, squid, arachnids and humans (Lewis & South, 2012; Albo *et al*., 2014; Lewis *et al*., 2014). Despite the ubiquity of this behaviour, little effort has been made to conceptualise the evolution of nuptial gift-giving within a general modelling framework (Lewis *et al*., 2014; Iwasa & Yamaguchi, 2022). Recent models describing the evolution of nuptial gift-giving have focused on co-evolution between male nuptial gift-giving and female propensity to remate, and evolutionarily stable nuptial gift sizes (Kamimura *et al*., 2021; Iwasa & Yamaguchi, 2022), but a general framework describing the conditions under which nuptial gift-giving can be initially favoured by selection is needed to understand when gift-giving should evolve.

Nuptial gift-giving may allow males to increase fitness by acquiring additional mates, indirect benefits (by increasing offspring fitness), prolonged copulations, and success in sperm competition (Albo *et al*., 2013; Ghislandi *et al*., 2014; Lewis *et al*., 2014). However, this potential fitness increase comes at the expense of producing a nuptial gift, which may be costly in terms of time and resources. Females may increase their fitness by receiving nutritionally valuable nuptial gifts, but expressing a preference for males with gifts might result in a mating opportunity cost if available males without gifts are rejected. With respect to nuptial gift-giving, the evolutionary interests of both sexes may not always fully overlap. This can cause sexual conflict, which is a difference in the fitness interests between sexes that occurs when an interaction results in a situation where individuals cannot both achieve an optimal outcome (Parker, 2006). For example, under some conditions, it might be optimal for males but not females to mate if females do not benefit from mating with males without nuptial gifts.

Much work has sought to explain how gift-giving tactics are maintained, with explanations including condition-dependent strategies, gift-giving as a way to decrease female aggression during copulation, or gifts as sensory traps (Lubin & Bilde, 2007; Toft & Albo, 2016; Ghislandi *et al*., 2018; Albo *et al*., 2019). An example of such a system is the nuptial gift-giving nursery-web spider *Pisaura mirabilis*, where males may court females with or without nuptial gifts (Bristowe & Locket, 1926; Tuni *et al*., 2013). Here, males may provide females with costly nuptial gifts in the form of captured arthropod prey, and females may exhibit preference for males with a nuptial gift by rejecting males without a nuptial gift (Albo *et al*., 2013).

We develop a general framework for investigating the evolution of nuptial gift-giving and choosiness using a time-in, time-out modelling approach and an individual-based model (Clutton-Brock & Parker, 1992). Specifically, we derive conditions under which selection will favour male search for nuptial gifts and female rejection of gift-less males. We show that selection for searching and choosiness depends on whether a threshold fitness value of the nuptial gift is exceeded. Our model demonstrates the importance of nuptial gift cost, sex ratio, and mate encounter rate in determining the threshold above which selection will favour the evolution of nuptial gift-giving. Importantly, we show that the threshold value differs for male searching and female choosiness. We complement the predictions of our analytical model by formulating an individual-based model, which further supports the main theoretical results. We apply our model to an example system with nuptial gifts, the nursery web spider *Pisaura mirabilis*, where we use experimental data to estimate a key model parameter. Our results provide a general framework for understanding why nuptial gift-giving evolves in some systems and not in others, how the evolution of nuptial gift-giving can give rise to sexual conflict, and it provides insight into the evolution of worthless and deceitful nuptial gifts, which occur in several different taxonomic groups (LeBas & Hockham, 2005; Ghislandi *et al*., 2014).

### Model

We use a time-in and time-out model (Clutton-Brock & Parker, 1992; Kokko & Monaghan, 2001; Kokko & Ots, 2006) in which choosy (female) and non-choosy (male) individuals spend some period of time within the mating pool searching for a mate (time-in) followed by a period outside the mating pool (time-out). During time-out, females spend some duration of time (*T*_f_) gestating or rearing (hereafter ‘processing’) offspring. We define the number of offspring produced by a female per reproductive cycle as *λ*. Since females enter time-out after mating, this is equivalent to assuming a system with sequential polyandry. For simplicity, we assume male time to replenish sperm is negligible, but males can spend some duration of time out of the mating pool searching for nuptial gifts. We define *T*_m_ as the time until a gift is found by a searching male.

#### Analytical model

The probability that a male who searches for an interval *T*_m_, and does not die during this period, finds a nuptial gift is given by,

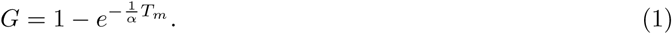

In Eq. 1, *α* defines expected search time before encountering a nuptial gift. Thus, the probability of finding a nuptial gift is higher the more time is spent searching. During time-in, individuals encounter conspecifics at a rate of *R*. A focal individual will therefore encounter conspecifics of the opposite sex at a rate of *R/*2 if the ratio of males to females in the mating pool (*β*) is equal. More generally, males will be encountered at a rate of *Rβ/*(*β* + 1), and females will be encountered at a rate of *R/*(*β* + 1). An example of how the structure of the time-in time-out model applies to a system with nuptial gift-giving is given in Figure 1. We assume that mating with a nuptial gift increases expected offspring production by an increment of *γ*.

**Figure 1:**
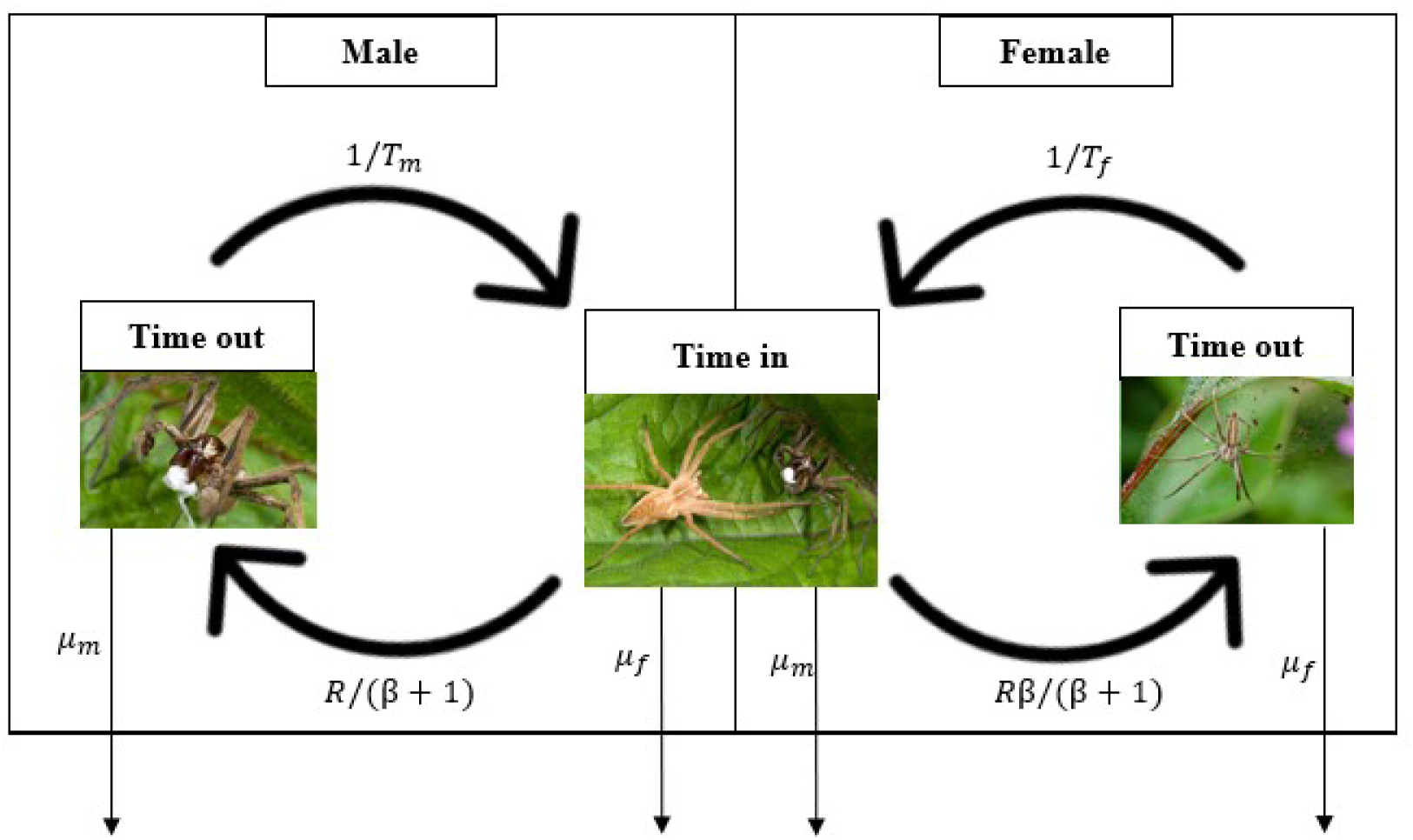
Conceptual figure inspired by Kokko and Ots (2006) illustrating how the modelling framework maps onto an example of a system wherein nuptial gifts are used, here *Pisaura mirabilis*. Males have a probability of obtaining a nuptial gift while in time-out, which will affect their probability of mating while in time-in. They return to the mating pool (time-in) at a rate determined by the time spent searching for a nuptial gifts (*T*_m_) and leave the mating pool (i.e. enter time-out) following the female encounter rate, which is dependent on the ratio of males to females (*β*) and the encounter rate (*R*). The choosy sex (females) enter the mating pool at a rate depending on the time spent processing offspring (*T*_f_) and leave the mating pool (i.e. enter time-out) at a rate that is dependent on *β* and *R*. Males and females undergo mortality *µ* during time-in and time-out. Image left to right: (1) male *P. mirabilis*. (2) male *P. mirabilis* presenting nuptial gift (white) to female. (3) Female *P. mirabilis* protecting offspring. Photos: Alamy.

Mortality of *µ*_f_ and *µ*_m_ for females and males, respectively, occurs both in and out of the mating pool. Following Kokko & Ots (2006), we assume *µ*_f_ = *µ*_m_ = 1. Assuming identical mortality both in and out of the mating pool means that individual phenotypes (search strategy and choosiness) do not affect mortality. This allows us to use a rate-maximisation model wherein maximisation of reproductive rate while alive is equivalent to maximising lifetime reproductive success (Kokko & Jennions, 2008; Jennions & Fromhage, 2017). Here maximisation of lifetime reproductive success is the same as maximising fitness because we do not model social interaction between relatives (Mylius & Diekmann, 1995; Fromhage *et al*., 2024). It should therefore be noted that if mortality differs inside and outside of the mating pool, reproductive rate while alive will no longer be equivalent to fitness. Nevertheless, we also investigated the evolution of both traits when mortality rates in and out of the mating pool differed substantially (see Supporting Information S1).

First, we find the fitness of males that search versus do not search for a nuptial gift. We then describe the fitness consequences of female choice to accept or reject males based on their provision of a nuptial gift. We can then calculate the thresholds *γ*_m_ and *γ*_f_ above which males and females are favoured by selection to search for nuptial gifts and exhibit choosiness for nuptial gifts, respectively.

##### Male fitness

During time-out, males can search for a nuptial gift. Males can adopt one of two strategies; either search or do not search for a nuptial gift. Males with the searching strategy continue to search until they find a nuptial gift, while males that do not search immediately re-enter the mating pool. Consequently, time searching for a nuptial gift will come at the cost of mating opportunities but might increase fecundity. We therefore need to model the expected length of time spent outside of the mating pool for males that search for nuptial gifts (*E*[*T*_m_]), which equals *α*. We can integrate search time *t* over the rate at which nuptial gifts are encountered (exp(*−*1*/α*)) to show *E*[*T*_m_] = *α*,

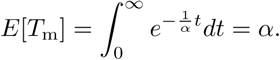

The rate at which a focal male that searches for a nuptial gift increases his fitness is therefore his reproductive output *λ*(1 + *γ*) divided by expected time spent searching for a nuptial gift (*α*) plus time spent in the mating pool, (*β* + 1)*/R*,

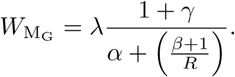

In contrast, a male that does not search for a nuptial gift has fewer offspring, but spends less time outside of the mating pool,

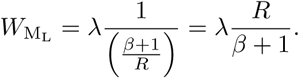

We can determine the conditions for which *W*_MG_ *> W*_ML_, isolating *γ* to find how large of a fitness benefit must be provided by the nuptial gift to make the search cost worthwhile, which simplifies to,

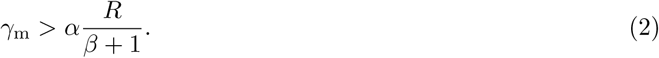

When this inequality holds, males are favoured to search until they find a nuptial gift, which would result in an average search time of *α*. If the male trait is instead continuous (i.e., males search for time period *T*_m_), it can be shown that the same threshold can be reached by evaluating the partial derivative of the male fitness function (Supporting Information S2). Hence, the thresholds are consistent under different assumptions concerning male searching strategy. Selection will cause males to search for nuptial gifts if the fitness increase to offspring exceeds the product of search time and female encounter rate.

##### Female fitness

During time-out, females process offspring over a duration of *T*_f_ (we assume that *T*_f_ *> α*, else females are not the choosy sex). When females re-enter the mating pool, they encounter males at a rate of *Rβ/*(*β* + 1). If a female encounters a male with a nuptial gift, we assume that she will mate with him. But if a female encounters a male with no nuptial gift, then she might accept or reject the male. If she rejects the male, then she will remain in the mating pool. The rate at which a female encounters a male with a nuptial gift is,

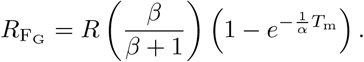

Expected time spent in the mating pool before a focal female encounters a male with a gift will be 1*/R*_FG_. The rate at which a female increases her fitness by being choosy and mating only when she encounters a male with a gift is,

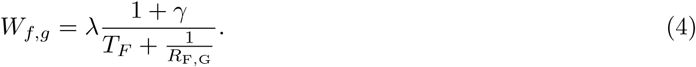

The top of the right-hand side of Eq. 4 gives her fitness increase, and the bottom gives the total time it takes to obtain this fitness. The *R*_FG_ is inverted because it represents the expected time to encountering a male with a gift. We can expand Eq. 6,

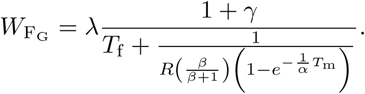

If the focal female is not choosy and accepts the first male that she encounters, then the rate at which she increases her fitness is,

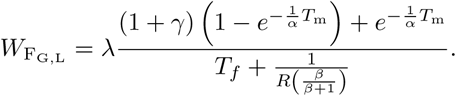

We then evaluate the conditions under which *W*_FG_ *> W*_FG,L_. We can isolate *γ* to determine how much fecundity must be increased to make choosiness beneficial (*γ*_f_),

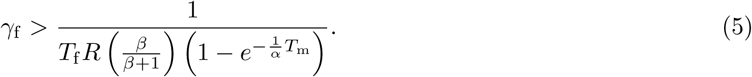

encounters males with nuptial gifts. Hence, female choosiness is ultimately determined by time spent out of the mating pool to process offspring (*T*_f_) and the rate at which a female in the mating pool encounters males with nuptial gifts.

##### Operational sex ratio

We assume that the sex ratio is equal upon maturation. Given this, Kokko & Monaghan (2001) show that the operational sex ratio depends on the probability of finding an individual in time-in,

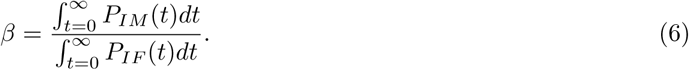

In Eq. 6, *P_IM_* (*t*) and *P_IF_* (*t*) are the probabilities of finding a male and female in time-in at time *t*, respectively. There is no closed form solution to the operational sex ratio, so we used recursion to calculate *β* values for a given *T*_f_, *T*_m_, and *R* (see Supporting Information S3),

#### Individual-based model

We formulate an individual-based time-in time-out simulation model to complement the predictions made by our analytical time-in time-out model above and further allow for coevolution between male search behaviour and female choosiness. Here we describe the details of initialisation, time-in (mating), time-out (reproduction and nuptial gift search), and mortality. We then summarise the simulations run and data collected.

##### Initialisation

Before the first time step, a population of *N* = 1000 individuals is initialised. Individuals are assigned to be female with a probability of 0.5, else male. Female offspring processing time for a focal individual *i* is fixed at *T^i^* = 2. Each individual is initialised with a diploid genotype that underlies female rejection probability of gift-less males (*ρ^i^*), and a diploid genotype that underlies male search time (*T_m_^i^*). Alleles can take any real value and combine additively to determine *ρ^i^* and *T_m_^i^*. For all simulations, initialised values are set to, *ρ^i^* = 0, and *T_m_^i^* = 0. All individuals are initialised outside of the mating pool in the first time step *t* = 1. The first time step then proceeds with females immediately entering the mating pool and males either entering the mating pool or searching for nuptial gifts.

##### Time-in

At the start of each time step, females and males in the mating pool remain in it. Females outside the mating pool enter the mating pool after processing offspring, and males enter it after searching for nuptial gifts (see ‘Time-out’ below). Up to Ψ = *Nψ* interactions between individuals can occur in a single time step, where *ψ* is a scaling parameter. In each time step, Ψ pairs of individuals are selected at random to interact. For each interaction, two different individuals are randomly selected from the population with equal probability. If the selected individuals are of different sexes, and both are in the mating pool, then a mating encounter occurs. If the male does not have a nuptial gift, then a focal female *i* will reject him with a probability of *ρ^i^*; if rejection occurs, then both individuals stay in the mating pool. If rejection does not occur, or the male has a nuptial gift in the mating encounter, then the individuals mate. Females then leave the mating pool and enter time-out to process offspring, and males leave and enter time-out to potentially search for new nuptial gifts (note that females and males might re-enter the mating pool immediately within the same time step given sufficiently low search time; see Time-out below).

##### Time-out

During time-out, offspring production and time outside of the mating pool are realised for each female by sampling from a Poisson distribution. A focal female *i* will produce Poisson(*λ*) offspring if no nuptial gift was provided or Poisson(*λ* + *γ*) if a gift was provided. Females remain outside of the mating pool to process offspring for Poisson(*T ^i^*) time steps. Offspring are added to the population immediately, with one *ρ^i^* and one *T_m_^i^* allele sampled from each parent with complete recombination. Mutation occurs independently for each allele with a probability of *ɛ* = 0.001, and if mutation occurs, then a new allele value is sampled from a normal distribution mean centred on the existing value with a standard deviation of *σ_ɛ_* = 0.1. Offspring sex is randomly assigned with equal probability as female or male. Female offspring are immediately placed in the mating pool, and male offspring are out of the mating pool to potentially search for nuptial gifts. After a female has spent Poisson(*T ^i^*) time steps outside the mating pool, she will re-enter it.

A focal male *i* outside the mating pool will enter it if he has searched for a fixed number of *T_m_^i^* time steps, which is also sampled randomly from a Poisson distribution, Poisson(*T_m_^i^*). If *T_m_^i^* = 0, then the male immediately returns to the mating pool (in the same time step). If *T_m_^i^* > 0, then the male must wait outside the mating pool for Poisson(*T_m_^i^*) time steps, but will enter the mating pool with a nuptial gift with a probability,

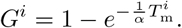

Males must always spend Poisson(*T_m_^i^*) time steps outside of the mating pool regardless of whether or not they are successful in obtaining a nuptial gift.

##### Mortality

At the end of each time step, mortality occurs first with a fixed probability for all adults in the population, then with a probability caused by carrying capacity *K* = 1000 applied to all individuals (adults and offspring). Mortality occurs in each time step with a fixed probability of *µ* regardless of the sex of the individual or its position in or out of the mating pool. If after this fixed mortality is applied, the total population size *N > K*, then individuals are removed at random with equal probability until *N* = *K*. Following adult mortality, a new time step begins with newly added offspring becoming adults.

##### Simulations

We ran simulations in which male search time and female choosiness evolved from an ancestral state of no searching and no choosiness. In all simulations, *N* was initialised at 1000 and *K* = 1000. Simulations ran for *t_max_* = 10000 time steps. We set *T*_f_ = 2, *ψ* = 3, and *λ* = 1 for all simulations, and we simulated across a range of *α* = *{*0.1, 0.2*, …,* 1.9, 2.0*}* and *γ* = *{*0, 0.1*, …,* 1.9, 2.0*}* parameter values for 5000 replicates. In a sensitivity analysis, we also re-ran simulations for values of *T*_f_ = 10 and *ψ* = *{*1, 2, 4, 6*}* (see Supporting Information S4), and simulated the evolution of male nuptial gift searching and female choosiness (fixing *T*_m_ = *α*) separately (see Supporting Information S4).

We additionally ran simulations in which males adopted one of two search strategies: either search in timeout until a gift is obtained, or do not search at all. This simulations were therefore more closely aligned with the assumptions of our analytical model, but results did not differ qualitatively (Supporting Information S5). The strategy adopted by a male in these simulations was based on an underlying trait for the probability of nuptial gift search. Like *ϕ^i^*, this trait was underpinned by a diploid genotype in which alleles had additive effects.

Summary statistics for mean trait values, population size, sex ratios, proportion of females and males in and out of the mating pool, and mean number of encounters per female and male within the mating pool were all calculated in the last time step. The C code used for simulating these IBMs also allows for the reporting of summary statistics in each time step. Additionally, it can simulate explicit space and individual movement through the landscape. A neutral evolving trait was also modelled to ensure that the code functioned as intended, and processes were compartmentalised into individual functions to facilitate code testing. All code is publicly available on GitHub (https://github.com/bradduthie/nuptial_gift_evolution).

##### Application to gift-giving spiders

We can produce an estimate of the fitness increment obtained by females when receiving a gift (*γ̂*) by using data on female *P. mirabilis* egg production and hatching success (Tuni *et al*., 2013). Tuni *et al*. (2013) found differences in egg production and hatching success in female *P. mirabilis* under different feeding regimes. Assuming these differences in feeding regimes correspond to eating versus not eating nuptial gifts, the mean number of offspring produced by a female who eats nuptial gifts can be calculated. We calculate these means using raw data from Tuni *et al*. (2013) and find that mean offspring produced by females receiving nuptial gifts is 25.74 *±* 0.96, while offspring produced by females without a nuptial gift is 6.00 *±* 2.1. Consequently, the relative gain in fitness from receiving a nuptial gift for a female is estimated to be *δ̂*_f_ = 25.74*/*6.00 = 4.29. Since the baseline fitness is 1, the increase in fitness resulting from a nuptial gift then becomes *γ̂* = *δ̂*_f_ *−* 1 = 3.29.

## Results

Our model predicts 4 zones that are delineated by inequalities 2 and 5 and describe the initial thresholds for the evolution of searching for nuptial gifts in males and choosiness for nuptial gifts in females. Figure 2a illustrates conditions under which no searching or choosiness evolves (zone A), only search (zone B) or choosiness (zone C) evolve, or both searching and choosiness evolve (zone D). Blue and red threshold lines in Figure 2 are defined by Eq. 2 and Eq. 5, respectively. Figure 2b illustrates the expansion of zone B as encounter rate decreases, which is consistent with the results of our IBM (Figure 3).

**Figure 2:**
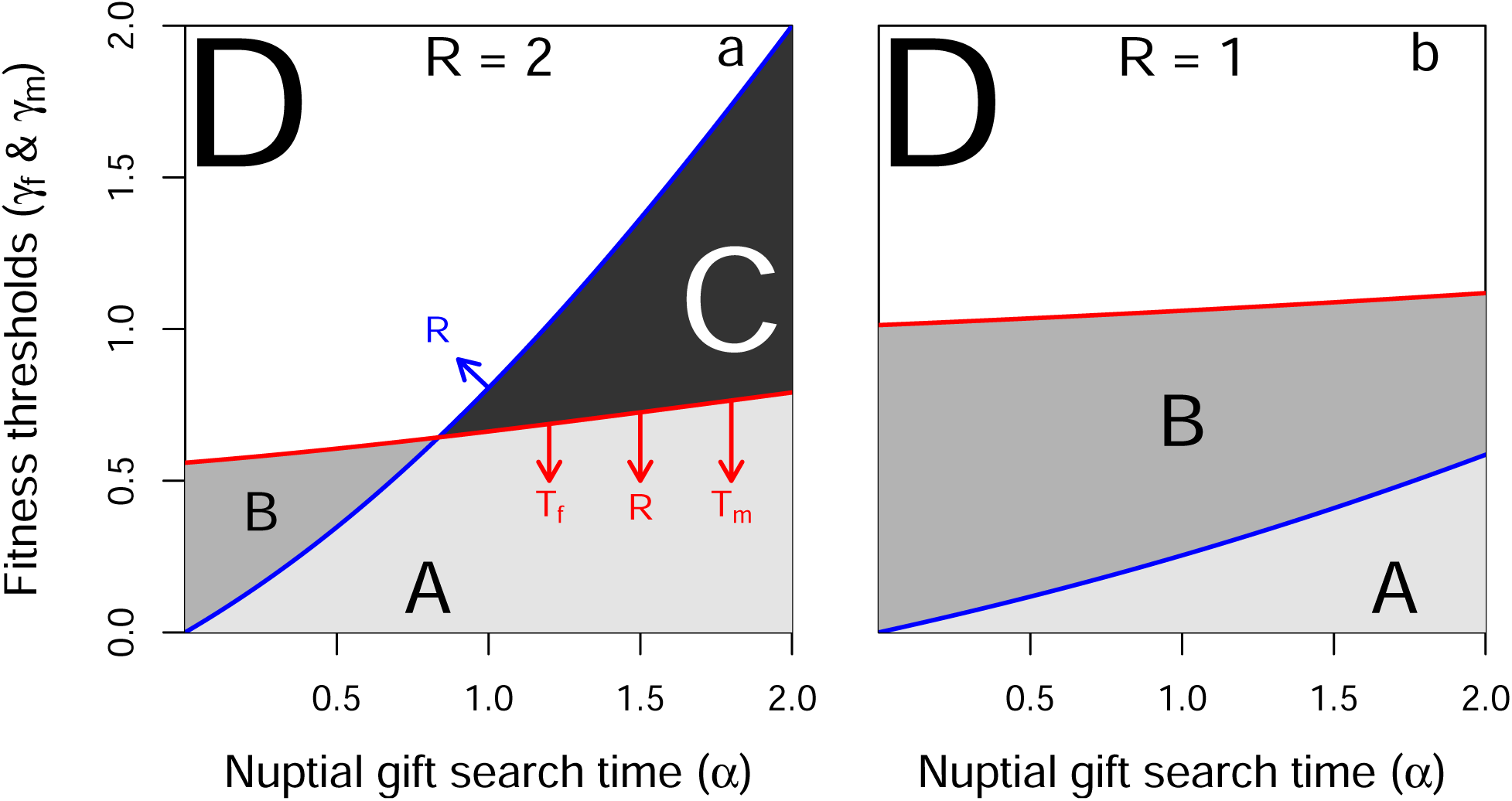
Fecundity increment above which males increase their fitness by searching for nuptial gifts (blue lines; Eq. 2) and females increase their fitness by rejecting males that do not offer gifts (red lines; Eq. 3). Parameter space includes areas in which males do not search for nuptial gifts and females are not choosy (A), males search but females are not choosy (B), females would be choosy but males do not search (C), and males search and females are choosy (D). Arrows in panel a indicate the effect of increasing encounter rate (*R*), female time-out (*T*_f_), and male search time (*T*_m_). As an example, curves for *T*_f_ = 2, and *T*_m_ = *α* are shown for values of *R* = 2 (panel a) and *R* = 1 (panel b). Females are assumed to be the choosy sex, which is maintained as long as *α < T*_f_.

**Figure 3:**
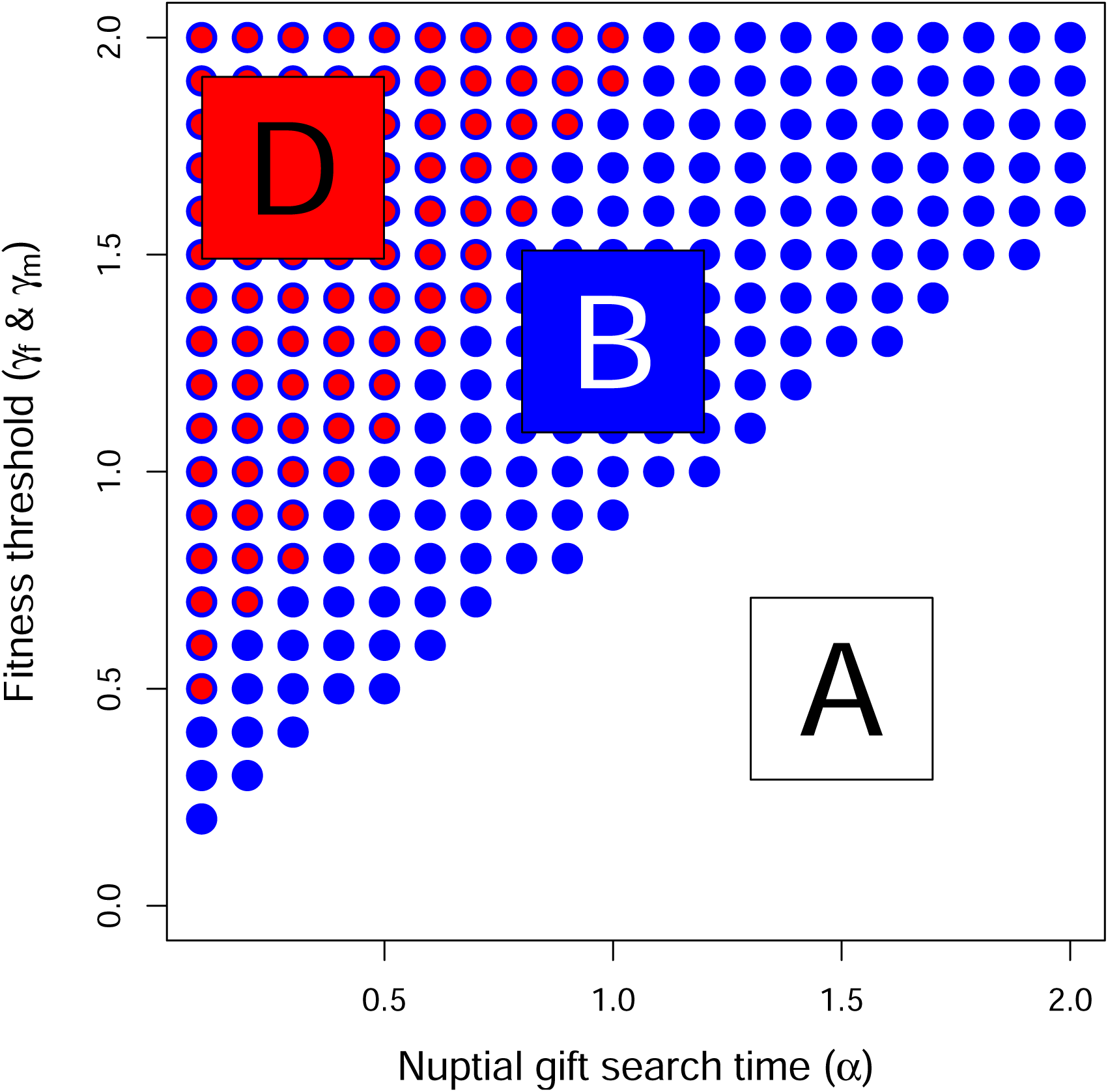
The coevolution of male search and female choosiness as a function of nuptial gift search time (*α*). Points show where the lower 95% confidence interval of female choosiness (red) and male search (blue) exceeds zero, indicating evolution of choosiness or nuptial gift search. Each point includes data from 3000 replicate simulations with identical starting conditions. Zones are identified that correspond to areas where males search and females are choosy (D), males search but females are not choosy (B), and males do not search and females are not choosy (A), as also depicted in Figure 2. The number of individuals in the population remained at or near carrying capacity of *K* = 1000. In each time step, up to 3000 total pair-wise interactions occurred. Expected female processing time was set to *T*_f_ = 2 time steps, and *γ* and *α* values in the range [0.0, 2.0] and [0.1, 2.0], respectively, were used.

For males, if nuptial gifts are not abundant and thus require a long time to find (i.e., high *α*), or if males encounter many females per unit time (i.e., high *R/*(1 + *β*)), then the nuptial gift must result in a high fitness increment for selection to favour searching. For females, if *γ* is sufficiently high, fitness is increased by rejecting males without gifts and mating only with males that provide nuptial gifts. As offspring processing time (*T*_f_), mate encounter rate (*Rβ/*(*β*+1)), or the probability of a male finding a nuptial gift (1*−*exp(*−T*_m_*/α*)) decrease, the fecundity increment above which selection will favour choosiness (*γ*_f_) increases. This can be understood intuitively by realising that rejecting a prospective male represents an opportunity cost for the female.

Individual-based model results recover zones A, B, and D (Figure 3). In other words, IBM simulations demonstrate that nuptial gift search in males, and choosiness in females, will evolve from an ancestral state of no searching and no choosiness in similar parameter space (Figure 3) as predicted by the analytical model (Figure 2b). Further, although it was not possible to parameterise our model for the *P. mirabilis* system in detail, our calculated value of *γ̂* = 3.29 is consistent with the evolution of nuptial gift searching and choosiness recovered by our analytical model and IBM.

## Discussion

Nuptial gift-giving has arisen several times independently throughout the animal kingdom (Lewis & South, 2012), so understanding how selection favours nuptial gift giving and choosiness is important for a broad range of mating systems. We provide a general framework that defines the necessary conditions for selection to favour the evolution of nuptial gift-giving. We show that males should give nuptial gifts if the value of a nuptial gift exceeds a threshold dependent on the encounter rate between females and males and the cost or time necessary to find or produce a nuptial gift. This result makes intuitive sense because if males rarely encounter females, time searching for a gift is a minor cost relative to mate search time. If males encounter many females, it is not worth seeking nuptial gifts unless gifts are very valuable since the male will meet many prospective mates, and nuptial gift search time might come at a cost of decreased mating opportunities. In practice, male biased sex ratios will not necessarily favour male search for nuptial gifts if the female encounter rate is very high, so the key variable is how often males and females encounter each other. If the search time or cost of finding a nuptial gift is high, nuptial gifts must be very valuable before search is favoured by selection. We also show that females should express choosiness for males with nuptial gifts if the value of the nuptial gift exceeds the inverse of the product of offspring processing time and encounter rate with males holding gifts.

### Threshold fitness values

We show that the threshold nuptial gift value at which females are favoured to express choosiness for nuptial gifts is rarely equivalent to the threshold value at which males are favoured to search for nuptial gifts, potentially leading to sexual conflict (Arnqvist & Rowe, 2005; Oliveira *et al*., 2008). Here, we are defining sexual conflict as occurring when interactions between sexes result in situation where both sexes cannot achieve an optimal outcome simultaneously (Parker, 2006). As an example, sizable areas of parameter space exists wherein the female optimum would be to exhibit preference for (and receive) nuptial gifts, while the male optimum is to not search for (and give) nuptial gifts (see Figure 2a, Zone C). This will lead to encounters where gift-less males would benefit the most from mating, but females would benefit most from mating and receiving a gift (i.e., evolutionary interests do not always overlap with respect to mating strategy between the sexes). In many systems, ecological variables such as search time required to find a nuptial gift will likely depend on prey abundance, which can vary substantially with time in some species with nuptial gift-giving (Ghislandi *et al*., 2018). Since several ecological variables likely affect the value of these thresholds, our results can be seen as providing some formalised description of why nuptial gift-giving only occurs in some but not all systems.

At first, the analytical model seems to suggests that nuptial gifts must cause a very high fitness increase (approximately 25%) before male search and female choosiness is favoured by selection (Figure 2). Similarly the IBM model seems to suggest that a fitness benefit of approximately 50% is required (see Figure 3). However, it is important to recognise that these thresholds depend on multiple parameters. For example, if female processing time (*T*_f_) is high, the female threshold for choosiness with respect to *γ* drops such that male search and female choosiness are favoured at lower *γ*. If *T*_f_ is sufficiently high, then an initially rare gift-giving trait might be favoured by selection even if the fitness benefit of a nuptial gift is low (assuming variation in choosiness for gifts existed among females). The effect that nuptial gifts have on fitness might vary across species, or even populations. Effects on female fecundity have been estimated in crickets, fireflies, butterflies, and spiders, but these estimates vary considerably. For example, one study of crickets (*Oecanthus nigricornis*) found that more nuptial feeding could increase female reproductive lifespan by approximately 50% (Brown, 1997), and a study on fireflies (*Photinus ignites* and *Ellychnia corrusca*) reported an increase in fecundity of 73% for females consuming multiple nuptial gifts (Rooney & Lewis, 2002). However, other studies on different systems have reported very varying effects or even no effect at all (Bergström & Wiklund, 2002; Rooney & Lewis, 2002; Maxwell & Prokop, 2018; Gao *et al*., 2019).

We modelled the evolution of nuptial gift-giving using both a mathematical model and an individual-based model. Our mathematical model makes simple assumptions about the relationship between nuptial gift search time (*α*), conspecific encounter rate (*R*), female processing time (*T*_f_), and the fitness increment of a nuptial gift for reproductive output (*γ*). It then derives the threshold *γ* values above which males increase their fitness by searching (*T*_m_ *>* 0) for a nuptial gift (*γ*_m_) and females increase their fitness by choosing to reject males without gifts (*γ*_f_). In contrast, our IBM models individuals over discrete time steps, and key processes of nuptial gift acquisition, conspecific encounters, and female processing are stochastic and varying among individuals. The mathematical model and IBM make qualitatively identical predictions (compare Figure 2b versus Figure 3), but differences between the two models inevitably lead to quantitative differences. For example, *γ*_f_ increased more rapidly with increasing *α* in the IBM compared to the analytical model. Some differences are expected to occur due to stochastic effects inherent to IBMs (e.g., Wilson *et al*., 2003). Other differences are more likely caused by more subtle assumptions between, and limitations of, the two models. In general, the IBM did not do a good job of controlling for conspecific interaction rate, making it difficult to directly compare *R* between models. The IBM also allowed for coevolution between male search and female choosiness (Figure 3), which was not allowed in the analytical model (Figure 2). It was not our goal to exactly recover the quantitative predictions of the analytical model in our IBM. Future development of the IBM could further bridge the gap between models while also developing new theory on how aspects of the system such as explicit space, individual life history, or genetics affect the evolution of nuptial gift-giving behaviour.

#### Nuptial gift-giving theory

When modelling nuptial gift evolution, the challenge is to construct a framework that captures the frequency-dependent selection between male nuptial gift-giving and female choosiness for nuptial gifts, and we do this using a time-in, time-out model. Recent studies have modelled some frequency-dependent aspect of nuptial gift giving using evolutionary game theory (Maynard Smith, 1982; Vincent & Brown, 2005). Two such studies formulated a quantitative genetics model to study evolutionarily stable nuptial gift sizes in populations where the female propensity to re-mate was evolving (Kamimura *et al*., 2021; Iwasa & Yamaguchi, 2022). The results obtained in these studies complement our results by giving equilibrium solutions to the evolutionary stable nuptial gift size, whereas we determine the general conditions under which nuptial gift-giving will evolve as given by the inequalities we derive.

Kokko & Mappes (2013) considered the evolution of female choosiness as a function of mating and mortality rates and found that choosiness may generally be a costly trait which is not always expected to evolve. Our results suggest that this is also the case when coevolution with the relevant male trait is taken into account. Numerous studies have modelled mate choice as a function of a single variable or effect (for review, see Edward, 2014), such as direct benefits, indirect benefits or so-called chase-away sexual selection (Holland & Rice, 1998; Kokko *et al*., 2003). Our results describe the evolution of mating choice from a different perspective by taking ecological variables and coevolution into account at the same time.

Other modelling frameworks have made general predictions about sexually selected traits, and these predictions are not mutually exclusive to those made by our model. For example, the good genes hypothesis predicts that costly traits such as nuptial gift-giving can be favoured since males enduring the cost of a nuptial gift signals to females that their genes confer high fitness precisely because they can afford this cost (Kirkpatrick, 1996; Byers & Waits, 2006; but see Fromhage & Henshaw, 2022). In other words, costly sexually selected traits are favoured because they are indicators of overall genetic quality (Martinossi-Allibert *et al*., 2019). Because of this, nuptial gift-giving could be a case of condition-dependence where engaging in nuptial gift-giving is only favourable for male in good condition (e.g., males capable of successful search (Maynard Smith, 1982; Engqvist & Taborsky, 2015; Ghislandi *et al*., 2018)). In general, our model demonstrates how nuptial gift-giving initially evolves before other mechanisms, such as good gene effects, become relevant.

A nuptial gift can also constitute a dishonest signal of good body condition since worthless, deceptive nuptial gifts have evolved in several systems (LeBas & Hockham, 2005; Ghislandi *et al*., 2014). This is also the case in *P. mirabilis* where males will wrap plant parts or an empty exoskeleton in silk, as opposed to an arthropod prey, and use this as a nuptial gift (Albo *et al*., 2011; Ghislandi *et al*., 2014). In such systems, worthless nuptial gifts have been shown to reduce the likelihood that a male is rejected by a female compared to the case where no nuptial gift is given. However, males offering worthless nuptial gifts may be at a slight disadvantage in sperm competition since worthless gifts result in a shorter copulation duration and hence less sperm transfer (Albo *et al*., 2013; Ghislandi *et al*., 2014). Worthless gifts should not result in any paternal care benefits to the male since the offspring he may sire will not gain nutrition from a worthless nuptial gift.

Given our modelling framework, worthless nuptial gifts may be expected to evolve in cases where females are discriminating in favour of nuptial gifts, but the cost of search time for a true nuptial gift is very high such that selection will not favour male search. This scenario would correspond to zone C of Figure 2 where the value of the nuptial gift exceeds the female fitness threshold for choosiness to be favoured, but due to high search time, selection will not favour male search for true nuptial gifts. Our model also predicts the possibility of the opposite scenario, in which males provide nuptial gifts, but females do not exhibit preference for nuptial gifts (zone B of Figure 2). Interestingly, an example of such system has been documented by a recent study of the genus *Trechaleoides*, which contains two species with true nuptial gift-giving, but a lack of preference for nuptial gifts among female (Martínez Villar *et al*., 2023).

The main drivers of male nuptial gift-giving are thought to be indirect fitness benefits and increased success in sperm competition, since providing a nuptial gift can result in longer copulation duration which is correlated with increased sperm transfer along with female cryptic choice promoting males who provide nuptial gifts (Albo *et al*., 2011, 2013). However, nuptial gifts might also function to modulate female aggression and prevent sexual cannibalism (Bilde *et al*., 2006). In some systems, such as *P. mirabilis*, males have been shown to reduce the risk of being cannibalised by the female after mating when offering a nuptial gift, such that the nuptial gift may result in a “shield effect”, protecting the male (Toft & Albo, 2016).

#### Empirical implications

For experimentally estimated value of *γ*, simulations showed evolution of nuptial gift searching in males and choosiness for nuptial gifts in females. The model thus predicts that *P. mirabilis* living under conditions with the estimated fitness value of nuptial gifts should exhibit both search for nuptial gifts and choosiness for males with nuptial gifts, and this is what is observed in empirical populations. Parameterising *γ* with data from experimental studies may only yield a rough approximation of the true *γ*. This is because the estimated value of *γ* is based on data from current populations (rather than ancestral populations, which are being simulated), and because the literature is inconclusive as to how much (if any) effect nuptial gifts have on female fitness (Maxwell & Prokop, 2018). The effect of nuptial gifts on female fecundity has been estimated in a variety of system such as crickets, fireflies, butterflies and spiders, but these estimates vary considerably between species suggesting a large positive effect to no effect at all (Bergström & Wiklund, 2002; Rooney & Lewis, 2002; Maxwell & Prokop, 2018; Gao *et al*., 2019).

Our model assumes sequential polyandry. That is, a system wherein female mating and reproduction with multiple males occurs in sequence, rather than multiple matings occurring before reproduction. In some systems with nuptial gift-giving, females have been documented to mate multiple times before reproduction occurs, including the genus of bark lice *Neotrogla* (Kamimura *et al*., 2021), and even our example system of *P. mirabilis* where females will sometimes engage in multiple mating before reproducing, especially if starved because multiple mating may result in more nuptial gifts (Toft & Albo, 2015; Matzke *et al*., 2022). It is unclear what effect (if any) assuming non-sequential polyandry would have on the threshold we derive. Under non-sequential polyandry, a viable strategy for females might be to accept any male (with or without gift) for fertilisation assurance, then exhibit a preference for nuptial gifts. This might make choosiness less costly since it would entail less of an opportunity cost to be choosy, and this could potentially make female preference for nuptial gifts more likely to evolve. Our model also assumes that males search in time-out, rather than contribute to parental care, which is likely to be accurate for most systems but not all. Expanding the model to explore these possibilities would be a worthwhile goal for future research.

## Conclusion

Overall, we found that a simple relationship between nuptial gift search time and mate encounter rate yields a threshold that determines whether selection will favour males that search for nuptial gifts. Similarly, we found that the threshold determining whether females will be favoured to reject males without nuptial gifts is also dependent on these variables, along with offspring processing time. Together, these thresholds describe the conditions under which nuptial gift-giving is expected to evolve. The applications of these thresholds are numerous. They can be used as a starting point for more complex or more system-specific models of nuptial gift-giving evolution. They can also provide novel insight into how populations can evolve to use worthless or token nuptial gifts.

## Author contributions

APC and ABD conceived the study. ABD constructed the modelling framework with input from APC. APC wrote the paper with input from ABD and ABD wrote the IBM model. TB and GB provided substantial comments on previous drafts and final text.

## Acknowledgements

Anders P. Charmouh was supported by the University of Aberdeen. A. Bradley Duthie was supported by the University of Stirling. Trine Bilde was supported by The Danish Council for Independent Research grant number 4002-00328B. We thank Maria Albo for making available data from a previous study which was useful for parameter estimating.

## Data availability

The simulation software was implemented in C and the full source code is available at https://github.com/bradduthie/nuptial_gift_evolution.

## Supporting Information

### S1: Sensitivity analysis of parameters in IBM

We investigated the sensitivity of our results to individual mortality during time-in (*µ*_in_) and time-out (*µ*_out_), female processing time (*T*_f_), and the potential for interactions between conspecifics (*ψ*). Across all of these simulations, there were challenges with statistical power. Evidence for the evolution of male search (blue points in figures) and female choosiness (red points in figures) was determined by the lower 95% bootstrapped confidence interval of *T*_m_ and *T*_f_ values being greater than zero, respectively. This required a lot of replicate simulations in the main text (Figure 3), especially for values just above predicted thresholds and for female choosiness. Computation time was a limiting factor, even using a compiled language (C) and with access to a computing cluster. Absence of points above threshold values are not necessarily evidence that evolution of male search or female choosiness is not predicted to evolve in these regions of parameter space, but it does indicate that evolution of these traits is not necessarily assured given the stochasticity inherent to the IBM. Additional simulations can be conducted using the C code in the ‘src’ folder of the GitHub repository (https://github.com/bradduthie/nuptial_gift_evolution). Below, we explain the parameter values used in the sensitivity analysis in more detail.

#### Mortality

We conducted a sensitivity analysis of the effect of the mortality parameters *µ*_in_ and *µ*_out_ (the probability of mortality in time-in and time-out, respectively, which we assumed to be equal for all individuals) on the evolution of male search and female choice using the IBM. The results revealed no correlation between the value of the mortality parameters and the evolution of male search or female choice (Fig. S2.1).

#### Female processing time

We also conducted a sensitivity analysis of female processing time *T*_f_. To do this, we ran simulations at default values, but with *T*_f_ = 10.0 (Figures S2.2).

#### Interactions between conspecifics

We conducted a sensitivity analysis on the encounter rate between conspecifics (*R*) by varying the value of our scaling parameter *ψ*. Under default simulations, *ψ* = 3. We also ran simulations in which *ψ* = 1 (Figures S2.3), *ψ* = 2 (Figures S3.4), *ψ* = 4 (Figures S3.5), and *ψ* = 6 (Figures S3.6), with all other parameters being set to default values.

**Figure S2.1:**
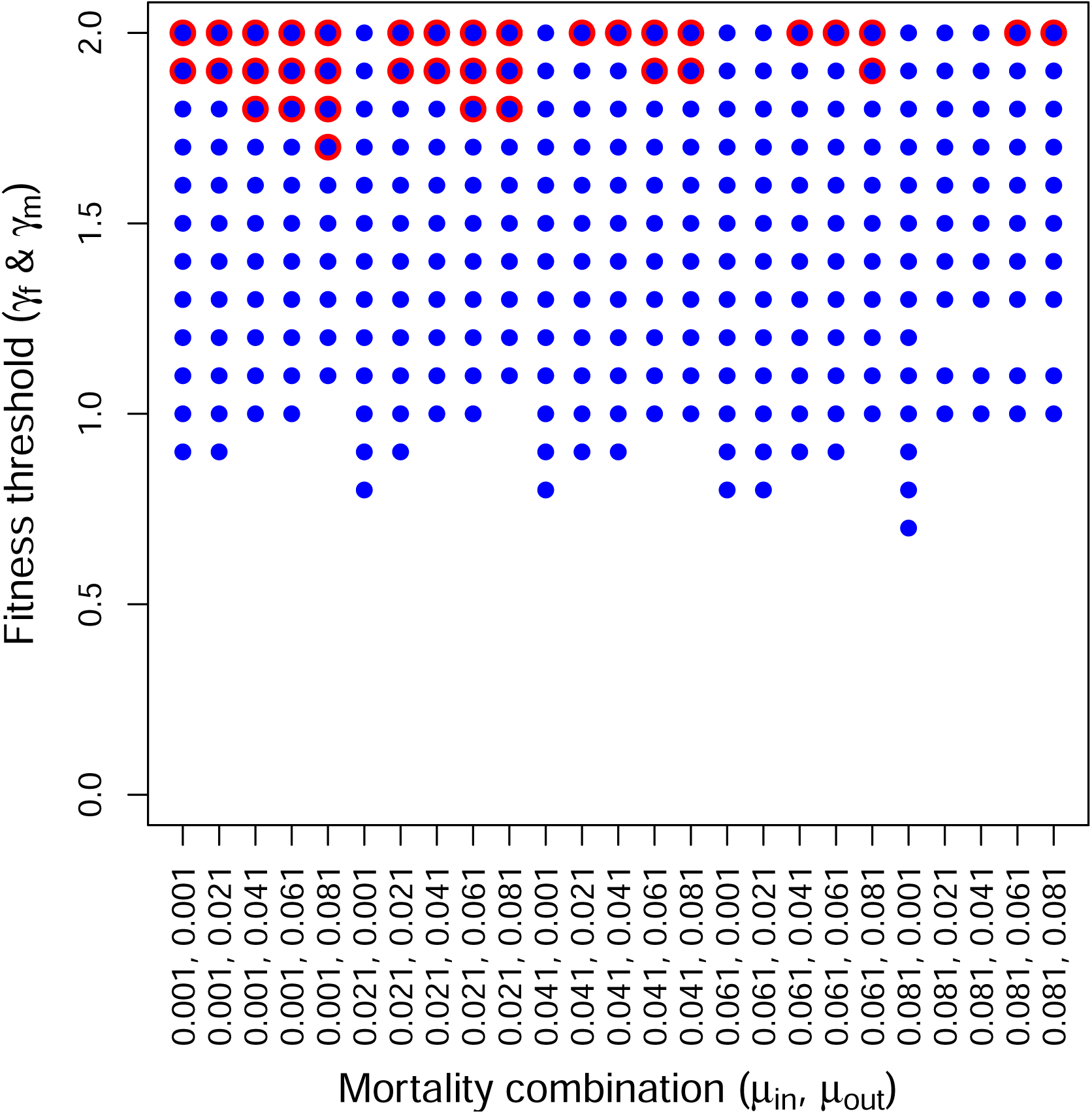
Evolution of both male search (blue) and female choice (red) under different combinations of the mortality rates *µ*_in_ and *µ*_out_ (mortality in time-in and out, respectively). The y-axis is the threshold fitness that leads to evolution of male search (blue) or female choice (red). The results show noise, but no correlation between the value of the mortality parameters and the propensity for male search and/or female choice to evolve. For each of the 25 *×* 20 combinations of *µ*_in_ and *µ*_out_, 3000 replicate simulations were run.

**Figure S2.3.**
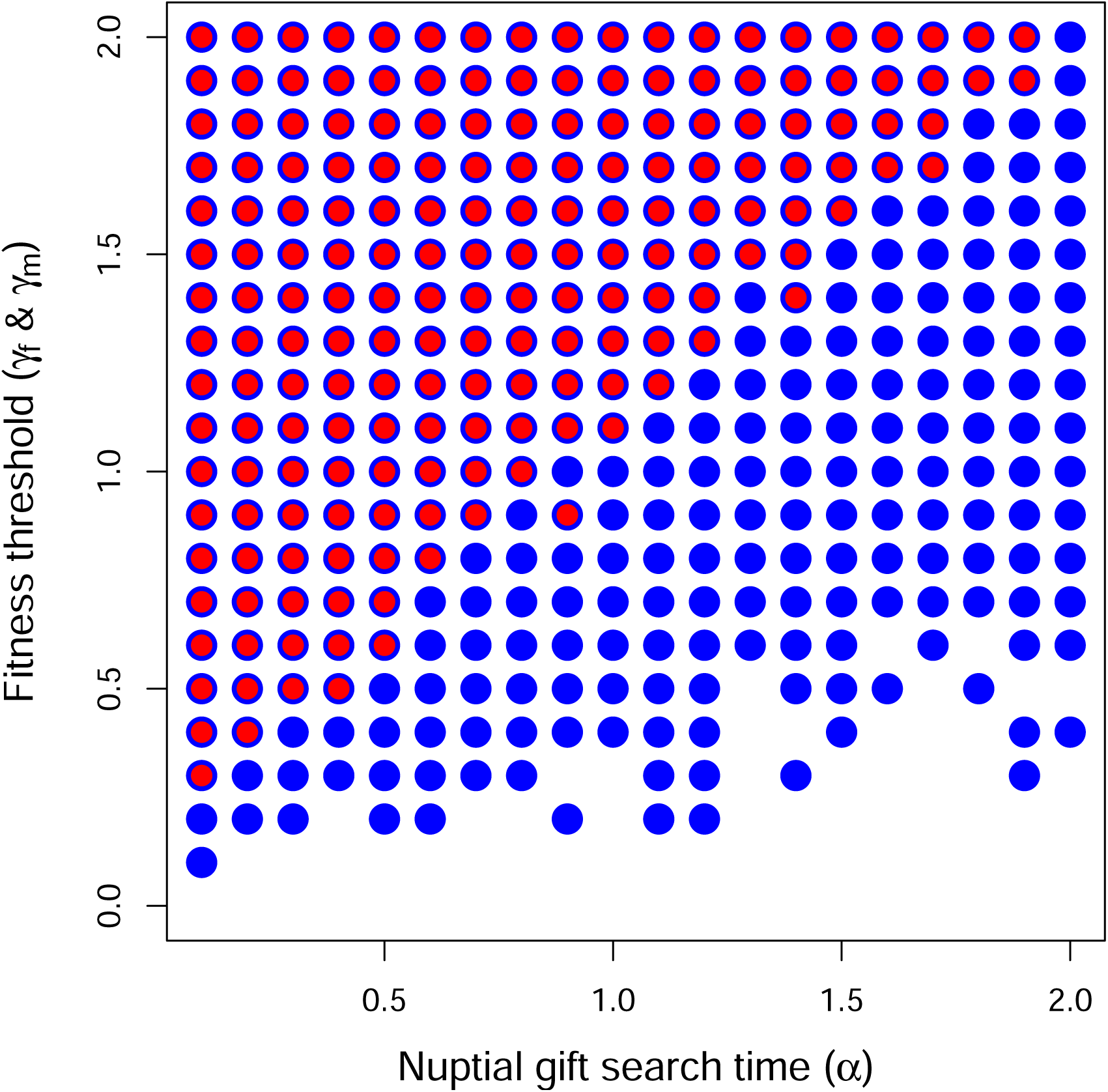
(*T*_f_ = 10.0): The coevolution of male search and female choosiness as a function of nuptial gift search time (*α*). Points show where the lower 95% confidence interval of female choosiness (red) and male search (blue) exceeds zero, indicating evolution of choosiness or nuptial gift search. Each point includes data from 3000 replicate simulations with identical starting conditions. Up to 3000 interactions occur between individuals in each time step (*ψ* = 3), potentially resulting in a mating interaction. The number of individuals in the population remained at or near carrying capacity of *K* = 1000. Expected female processing time was set to *T*_f_ = 10.0 time steps, and *γ* and *α* values in the range [0.1, 2.0] and [0.0, 2.0], respectively, were used.

**Figure S2.4.**
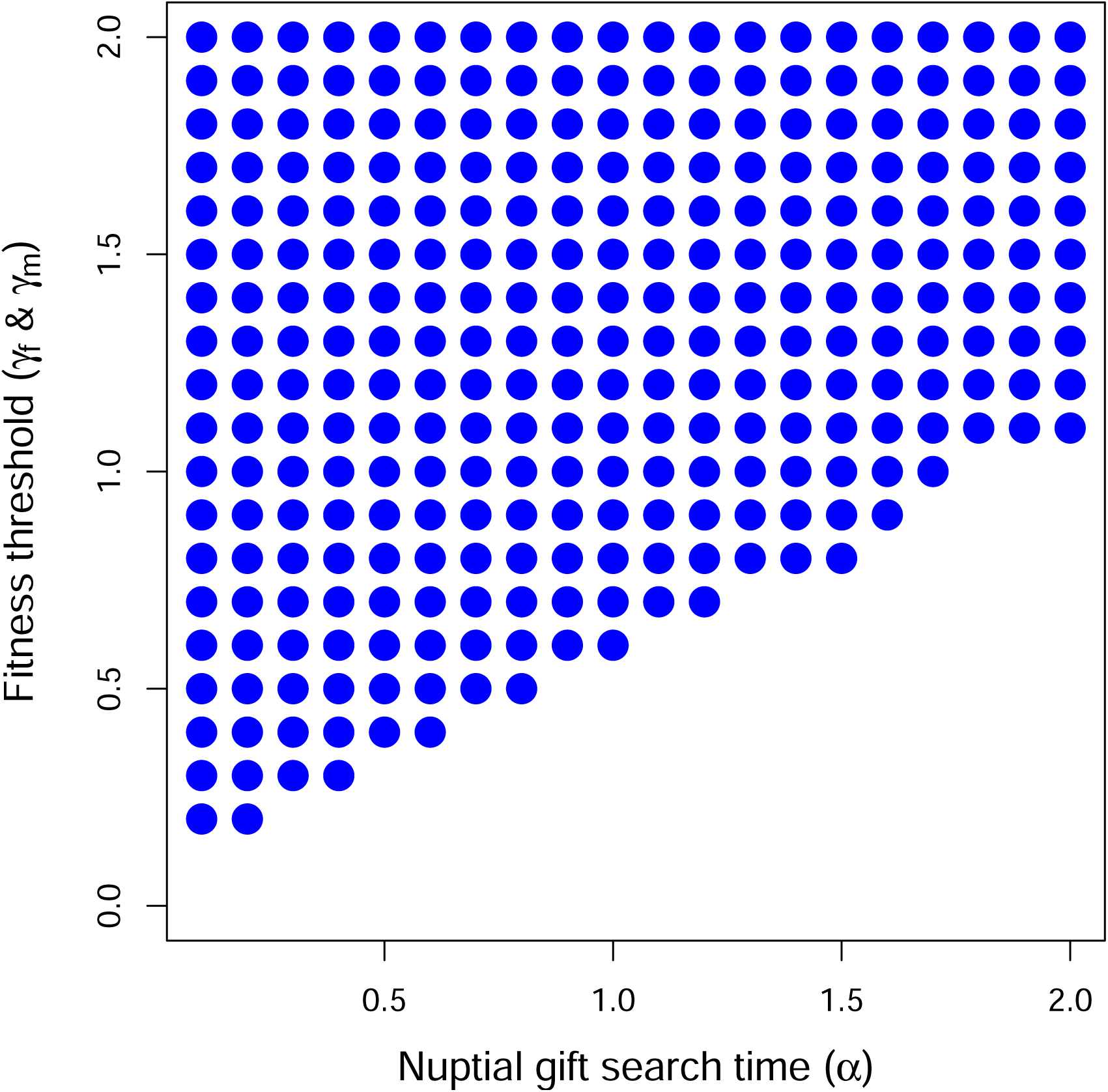
(*ψ* = 1): The coevolution of male search and female choosiness as a function of nuptial gift search time (*α*). Points show where the lower 95% confidence interval of where male search (blue) exceeds zero, indicating evolution of choosiness or nuptial gift search. Each point includes data from 3000 replicate simulations with identical starting conditions. Up to 1000 interactions occur between individuals in each time step (*ψ* = 1), potentially resulting in a mating interaction. The number of individuals in the population remained at or near carrying capacity of *K* = 1000. Expected female processing time was set to *T*_f_ = 2 time steps, and *γ* and *α* values in the range [0.0, 2.0] and [0.1, 2.0], respectively, were used.

**Figure S2.5.**
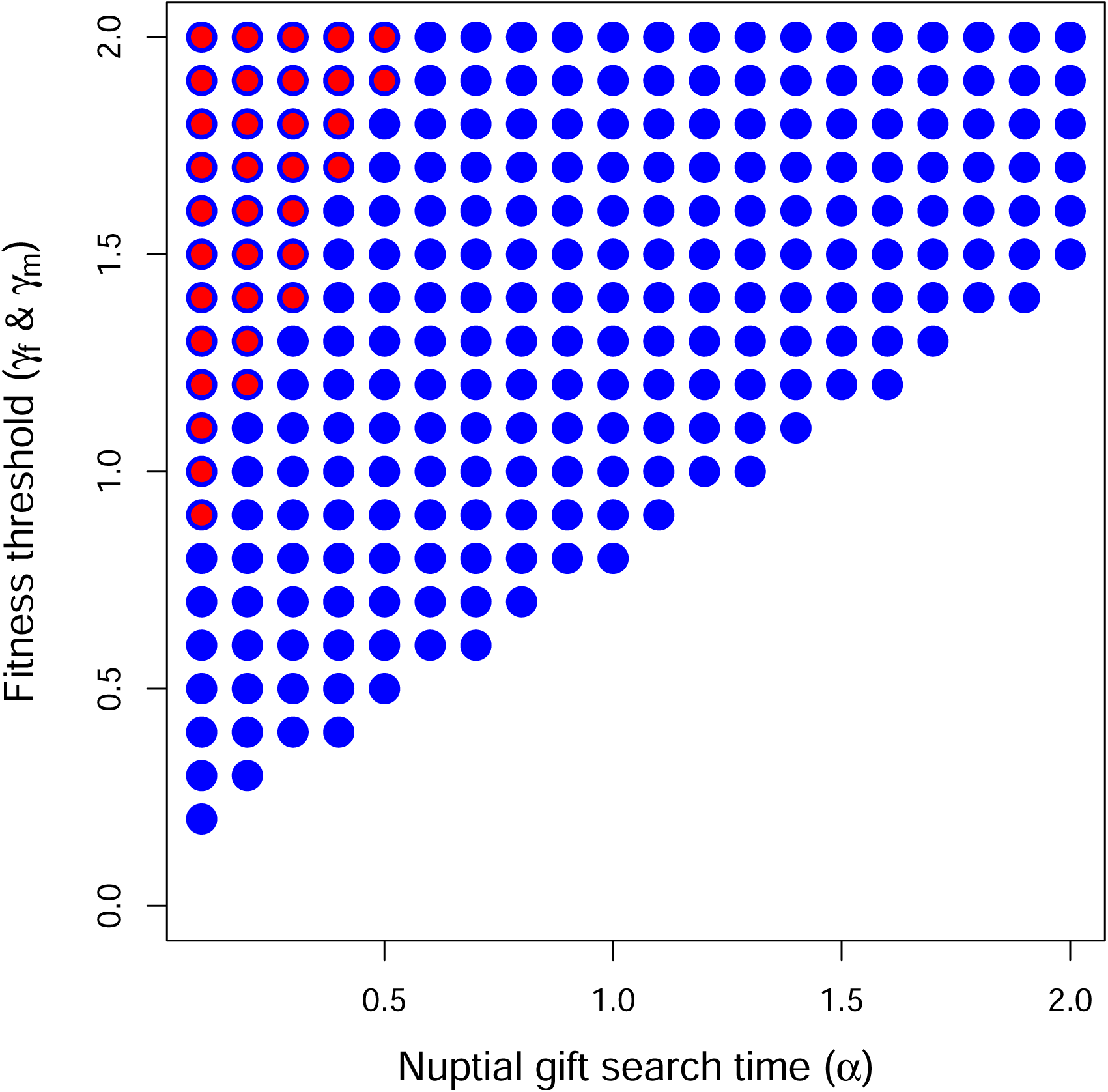
(*ψ* = 2): The coevolution of male search and female choosiness as a function of nuptial gift search time (*α*). Points show where the lower 95% confidence interval of where male search (blue) exceeds zero, indicating evolution of choosiness or nuptial gift search. Each point includes data from 3000 replicate simulations with identical starting conditions. Up to 2000 interactions occur between individuals in each time step (*ψ* = 2), potentially resulting in a mating interaction. The number of individuals in the population remained at or near carrying capacity of *K* = 1000. Expected female processing time was set to *T*_f_ = 2 time steps, and *γ* and *α* values in the range [0.0, 2.0] and [0.1, 2.0], respectively, were used.

**Figure S2.6.**
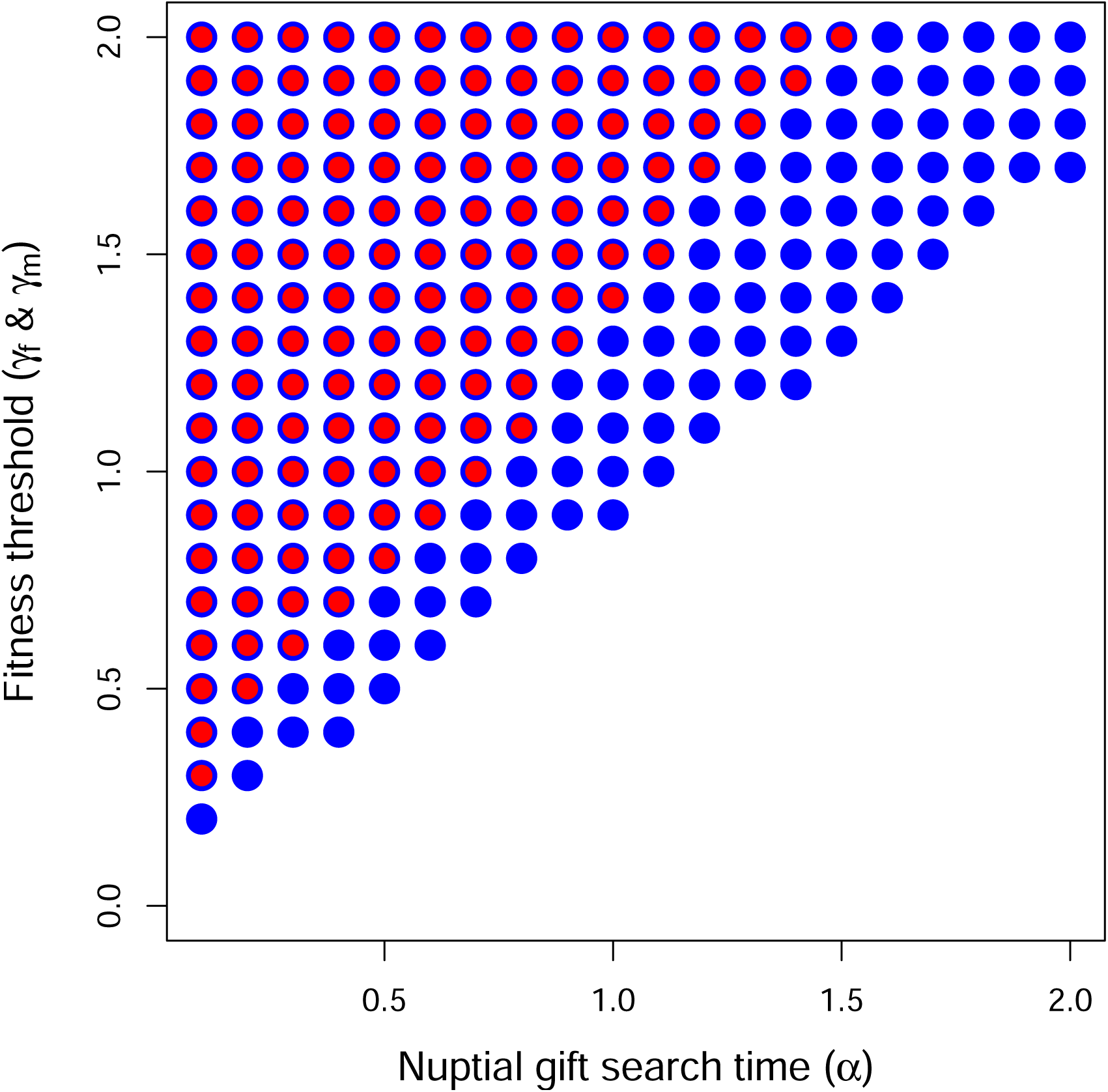
(*ψ* = 4): The coevolution of male search and female choosiness as a function of nuptial gift search time (*α*). Points show where the lower 95% confidence interval of where male search (blue) exceeds zero, indicating evolution of choosiness or nuptial gift search. Each point includes data from 3000 replicate simulations with identical starting conditions. Up to 4000 interactions occur between individuals in each time step, potentially resulting in a mating interaction (*ψ* = 4). The number of individuals in the population remained at or near carrying capacity of *K* = 1000. Expected female processing time was set to *T*_f_ = 2 time steps, and *γ* and *α* values in the range [0.0, 2.0] and [0.1, 2.0], respectively, were used.

**Figure S2.7.**
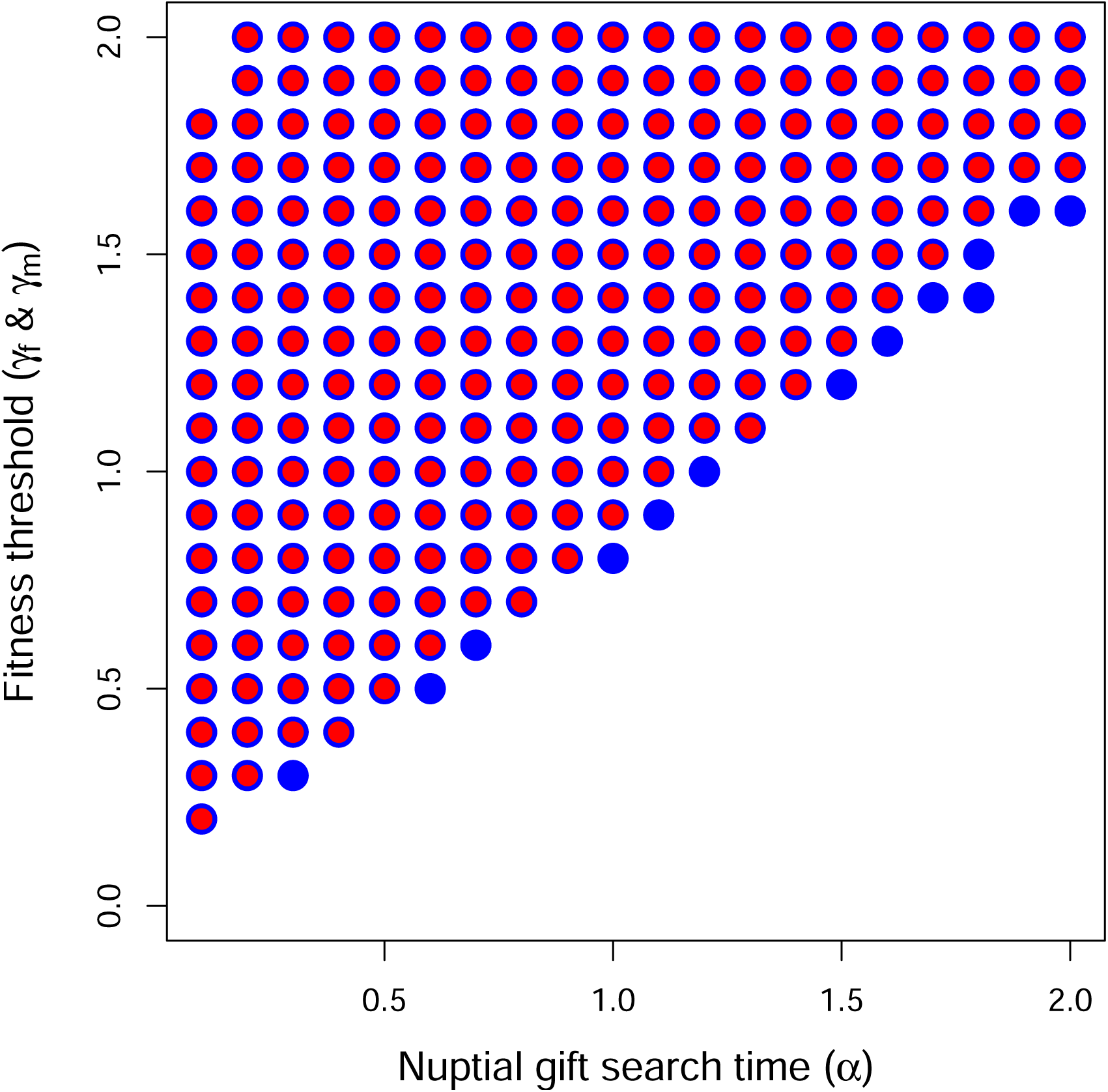
(*ψ* = 6): The coevolution of male search and female choosiness as a function of nuptial gift search time (*α*). Points show where the lower 95% confidence interval of where male search (blue) exceeds zero, indicating evolution of choosiness or nuptial gift search. Each point includes data from 3000 replicate simulations with identical starting conditions. Up to 6000 interactions occur between individuals in each time step, potentially resulting in a mating interaction (*ψ* = 6). The number of individuals in the population remained at or near carrying capacity of *K* = 1000. Expected female processing time was set to *T*_f_ = 2 time steps, and *γ* and *α* values in the range [0.0, 2.0] and [0.1, 2.0], respectively, were used.

### S2: Alternative derivation of male fitness threshold

In the main text, we assumed that males made the decision to search or not search for a nuptial gift. The expected length of time for which searching males are expected to remain outside of the mating pool is *E*[*T*_m_] = *α* (see Model). Alternatively, we can assume that males search for a pre-determined period of *T*_m_ and spend this full duration of *T*_m_ in the time-out phase, even if they succeed in finding a nuptial gift (as we assumed in our IBM). The probability that a male obtains a nuptial gift during this time is modelled in Eq. 1,

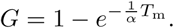

In Eq. 1, *α* is the amount of time expected to pass before a male encounters a nuptial gift. We assume that a male will only enter the mating pool with no gift if they are unsuccessful in obtaining a gift, so the probability that a male obtains no gift after *T*_m_ is,

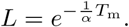

We assume that the fitness associated with receiving a nuptial gift versus no nuptial gift are *λ*(1 + *γ*) and *λ*, respectively. The rate at which males increase their fitness can then be defined as the expected fitness increase from their nuptial gift search divided by *T*_m_ plus the time spent in the mating pool waiting to encounter a mate,

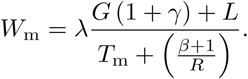

Our objective now is to determine the conditions under which a focal male increases his fitness by searching for a nuptial gift (*T*_m_ *>* 0) in a population of resident males that do not search (*T*_m_ = 0). Females are assumed to exhibit no choosiness for males with versus without nuptial gifts. Under such conditions, male fitness cannot be affected by female choice, so selection to increase *T*_m_ *>* 0 must be based solely on *α*, *β*, *R*, and *γ*.

To determine under what conditions male inclusive fitness increases with nuptial gift search time, we can differentiate *W*_m_ with respect to *T*_m_,

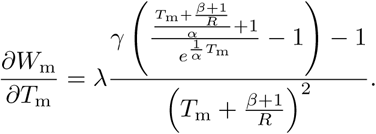

Because *T*_m_ = 0, the above simplifies,

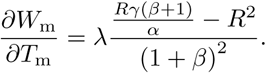

We can re-arrange the above,

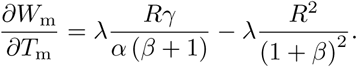

Note that if *R* = 0 or *λ* = 0, then, trivially, no change in fitness occurs (since females and males cannot mate or do not produce offspring). Fitness is increased by searching for nuptial gifts when *γ* is high, scaled by the expect search time needed to find a nuptial gift. A second term on the right-hand side is subtracted, which reflects a loss in fitness proportional to the encounter rate of potential mates in the mating pool. The threshold for which male inclusive fitness is not affected by searching for a nuptial gift are found by setting *∂W*_m_*/∂T*_m_ = 0 and solving for *γ* to recover Eq. 2 from the main text,

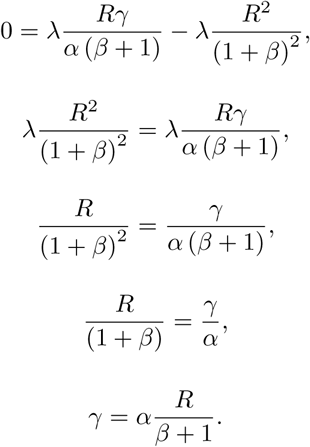

### S3: Operational sex ratio

We assume that the ratio of males to females is the same upon individual maturation. Consequently, the operational sex ratio *β* will be a function of *R*, *T*_f_, and *T*_m_ because these parameters determine the density of females and males in the mating pool versus outside of the mating pool. We start with the definition of *β* as being the probability of finding an individual in time-in (Kokko & Monaghan, 2001),

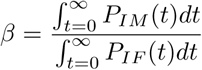

We can substitute the equations for *P_IM_* (*t*) and *P_IF_* (*t*), which define the probabilities of males and females being within the mating pool at time *t*, respectively.

We can therefore calculate *β* as below,

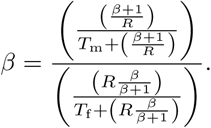

This can be simplified,

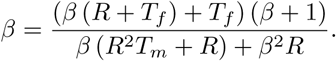

There is no closed form solution for *β*, but a recursive algorithm can be used to calculate *β* to an arbitrary degree of precision.

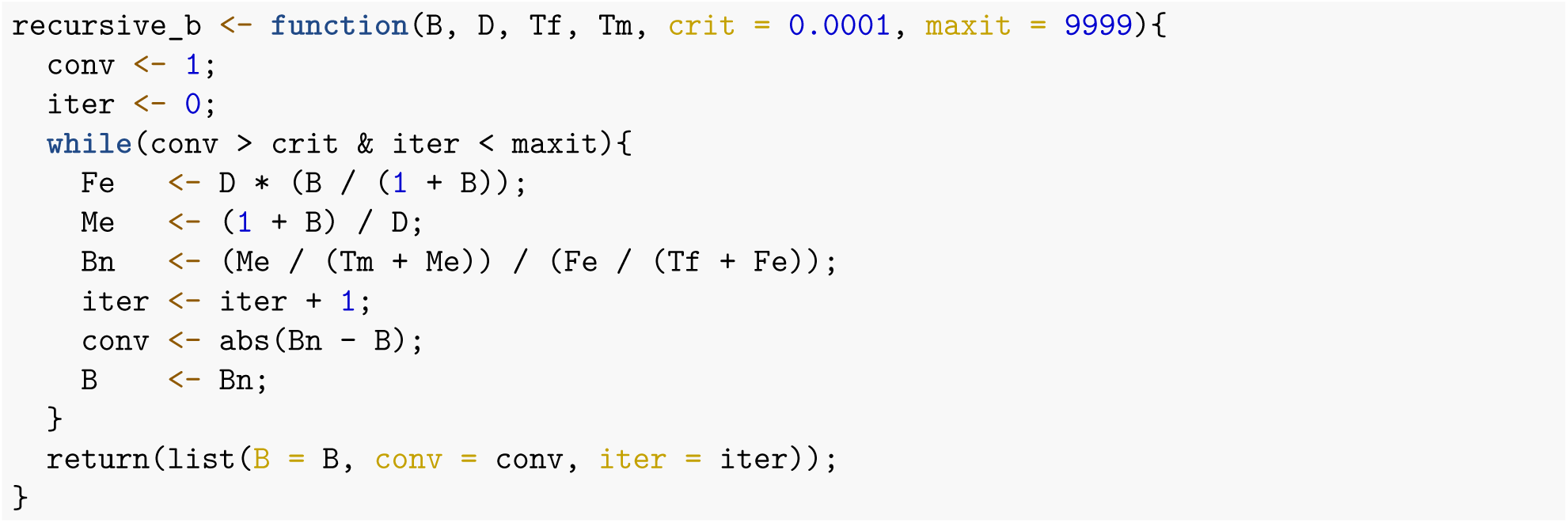

We used the above function to calculate values of *β* for the analytical model.

### S4: Separate evolution of male search and female choice

We used the individual-based simulation model to unpack the effect of coevolution on the evolution of male search and female choice. Here we replicated the simulations shown in the main text under the condition where only one trait at a time was allowed to evolve and studied how this affected the trait evolution.

First, we submitted a set of simulations wherein male search did not evolve, but was fixed at different values. Next, we ran the same set of simulations wherein male search evolved, but female choice was not possible. The results thus show how each trait evolves in the absence of any coevolution (Fig. S4.1).

**Figure S4.1:**
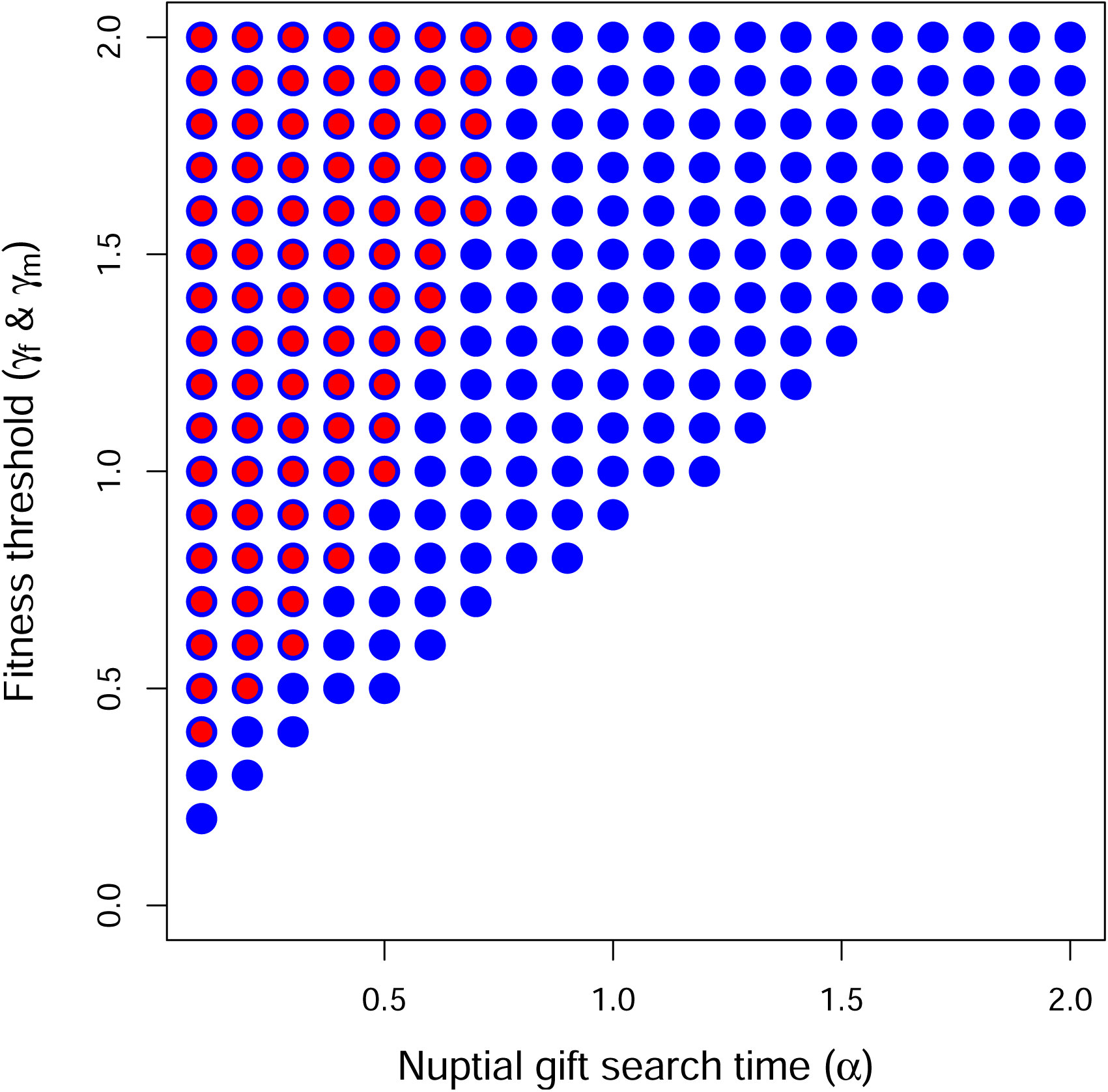
The separate evolution of male search and female choosiness as a function of nuptial gift search time. Points show where the lower 95% confidence interval of male search (blue) and female choosiness (red) exceeds zero, indicating evolution of nuptial gift search or choosiness. Each point includes data from 2 *×* 3000 replicate simulations with identical starting conditions. In the first batch, male search was constant and initialized at *T*_m_ = *α*, and female choice was evolving. In the second batch, male search was evolving, and there was no option for female choice. The parameters *T*_f_ = 2, and *γ* and *α* values were set within the range [0.1, 2.0] and [0.0, 2.0], respectively.

### S5: Male probability of search trait

In the individual-based model of the main text, a focal male *i* searches for a nuptial gift for a fixed number of Poisson(*T_m_^i^*) time steps. The male will remain outside the mating pool for this number of time steps regardless of whether or not he successfully obtains a nuptial gift. Here we instead define male search for nuptial gift as an all-or-nothing strategy to align with our analytical model. Males either search in time-out until they find a nuptial gift, or they do not search at all. In these simulations, *T_m_^i^* is instead defined as the probability that a focal male *i* searches for a nuptial gift (in the same way that *ρ^i^* is defined as the probability that a focal female *i* rejects a male without a nuptial gift). For males that decide to search, the time taken until a nuptial gift is obtained is randomly sampled from an exponential distribution with a rate parameter of *α*. This sampling time is then rounded to the nearest integer to determine the number of time steps that a focal male spends outside the mating pool before obtaining a nuptial gift.

Results are qualitatively identical to those in Figure 3 of the main text (Figures S5.1)

**Figure S5.1:**
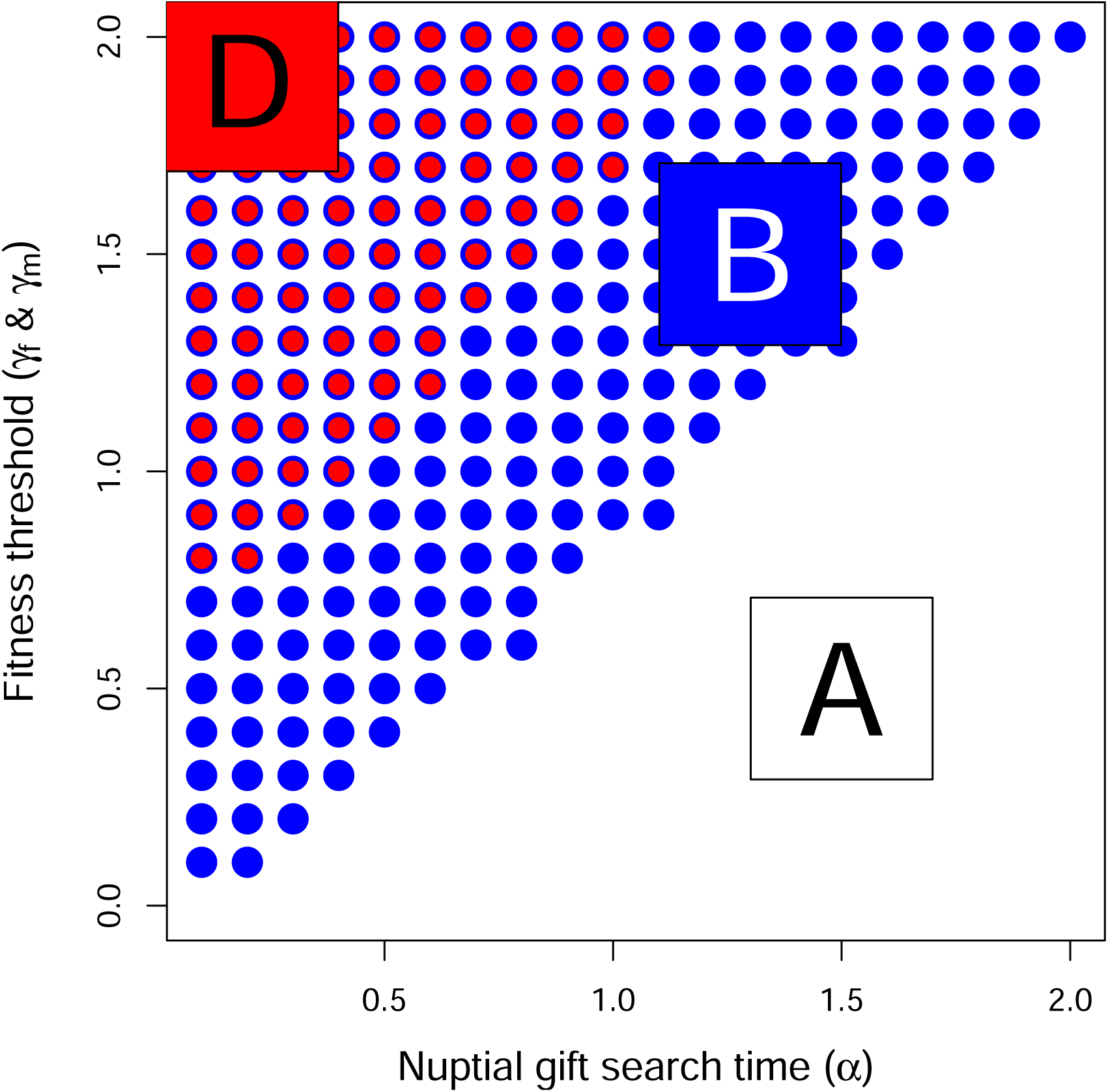
Coevolution of male search and female choosiness as a function of nuptial gift search time (*α*) when male strategy is binary: search or do not search. Points show where the lower 95% confidence interval of male search (blue) and female choosiness (red) exceeds zero, indicating evolution of nuptial gift search or choosiness. Each point includes data from 1000 replicate simulations with identical starting conditions. Zones are identified that correspond to areas where males search and females are choosy (D), males search but females are not choosy (B), and males do not search and females are not choosy (A), as also depicted in Figures 2 and 3 in the main text. The number of individuals in the population remained at or near carrying capacity of *K* = 1000. In each time step, up to 3000 total pair-wise interactions occurred. Expected female processing time was set to *T*_f_ = 2 time steps, and *γ* and *α* values in the range [0.0, 2.0] and [0.1, 2.0], respectively, were used.

