## Supporting Information for "A general model for the evolution of nuptial gift-giving"

Author list (anonymised)

[1] Author institutions and email

### Table of Contents

| Information | Page |
|---|---|
| S1: Sensitivity analysis of parameters in IBM | 2 |
| S2: Alternative derivation of male fitness threshold | 9 |
| S3: Operational sex ratio | 11 |
| S4: Separate evolution of male search and female choice | 12 |
| S5: Male probability of search trait | 13 |
| Reference | 15 |

### S1: Sensitivity analysis of parameters in IBM

We investigated the sensitivity of our results to individual mortality during time-in ($\mu_{\mathrm{in}}$) and time-out ($\mu_{\mathrm{out}}$), female processing time ($T_{\mathrm{f}}$), and the potential for interactions between conspecifics ($\psi$). Across all of these simulations, there were challenges with statistical power. Evidence for the evolution of male search (blue points in figures) and female choosiness (red points in figures) was determined by the lower 95% bootstrapped confidence interval of $T_{\mathrm{m}}$ and $T_{\mathrm{f}}$ values being greater than zero, respectively. This required a lot of replicate simulations in the main text (Figure 3), especially for values just above predicted thresholds and for female choosiness. Computation time was a limiting factor, even using a compiled language (C) and with access to a computing cluster. Absence of points above threshold values are not necessarily evidence that evolution of male search or female choosiness is not predicted to evolve in these regions of parameter space, but it does indicate that evolution of these traits is not necessarily assured given the stochasticity inherent to the IBM. Additional simulations can be conducted using the C code in the 'src' folder of the GitHub repository (https://github.com/AUTHOR_NAME/REPOSITORY_NAME). Below, we explain the parameter values used in the sensitivity analysis in more detail.

*Mortality*

We conducted a sensitivity analysis of the effect of the mortality parameters $\mu_{\mathrm{in}}$ and $\mu_{\mathrm{out}}$ (the probability of mortality in time-in and time-out, respectively, which we assumed to be equal for all individuals) on the evolution of male search and female choice using the IBM. The results revealed no correlation between the value of the mortality parameters and the evolution of male search or female choice (Fig. S2.1).

*Female processing time*

We also conducted a sensitivity analysis of female processing time $T_{\mathrm{f}}$. To do this, we ran simulations at default values, but with $T_{\mathrm{f}} = 10.0$ (Figures S2.2).

*Interactions between conspecifics*

We conducted a sensitivity analysis on the encounter rate between conspecifics ($R$) by varying the value of our scaling parameter $\psi$. Under default simulations, $\psi = 3$. We also ran simulations in which $\psi = 1$ (Figure S2.3), $\psi = 2$ (Figure S3.4), $\psi = 4$ (Figure S3.5), and $\psi = 6$ (Figure S3.6), with all other parameters being set to default values.

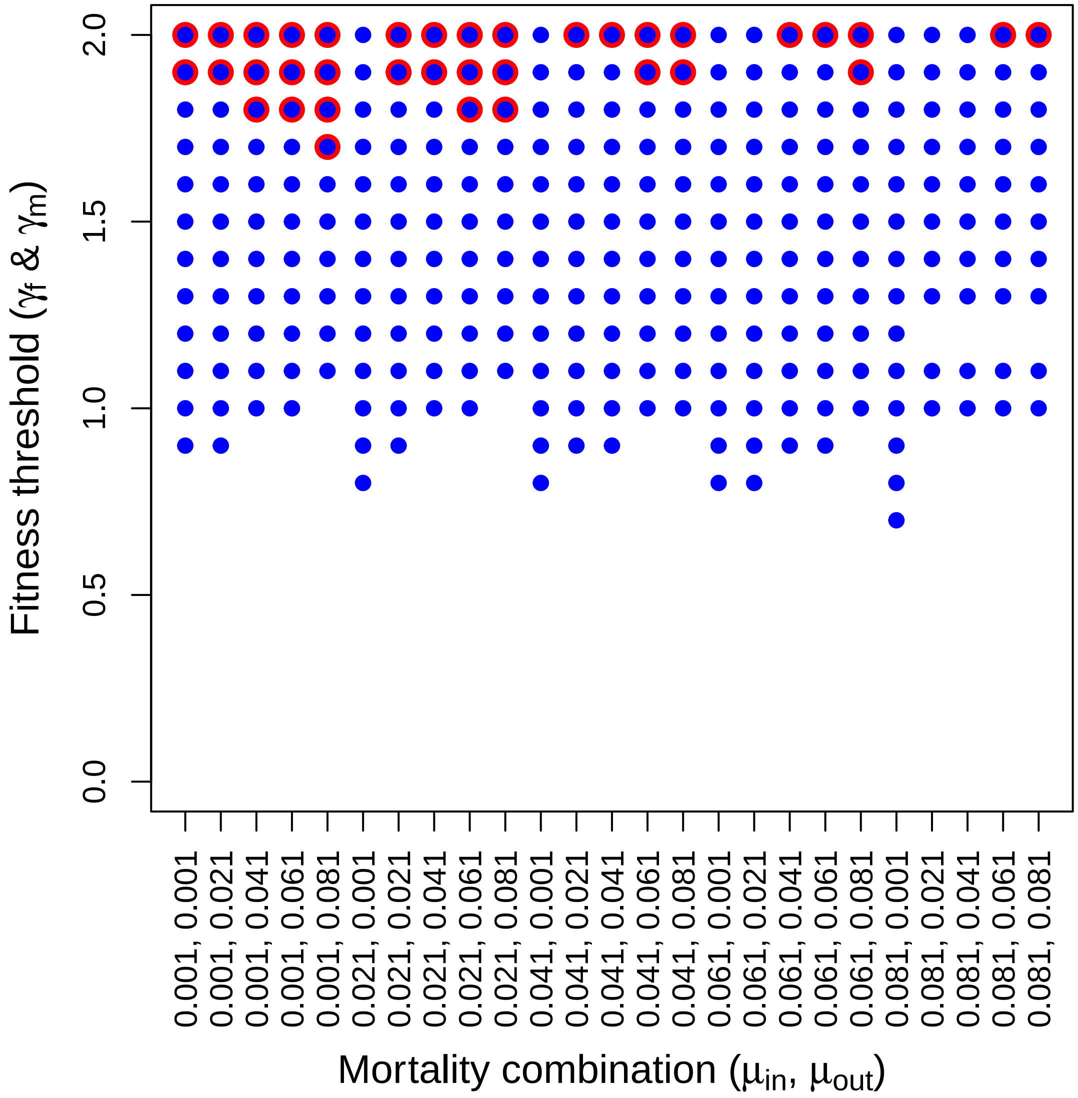


Figure S2.1: Evolution of both male search (blue) and female choice (red) under different combinations of the mortality rates $\mu_{\text{in}}$ and $\mu_{\text{out}}$ (mortality in time-in and out, respectively). The y-axis is the threshold fitness that leads to evolution of male search (blue) or female choice (red). The results show noise, but no correlation between the value of the mortality parameters and the propensity for male search and/or female choice to evolve. For each of the 25 × 20 combinations of $\mu_{\text{in}}$ and $\mu_{\text{out}}$, 3000 replicate simulations were run.

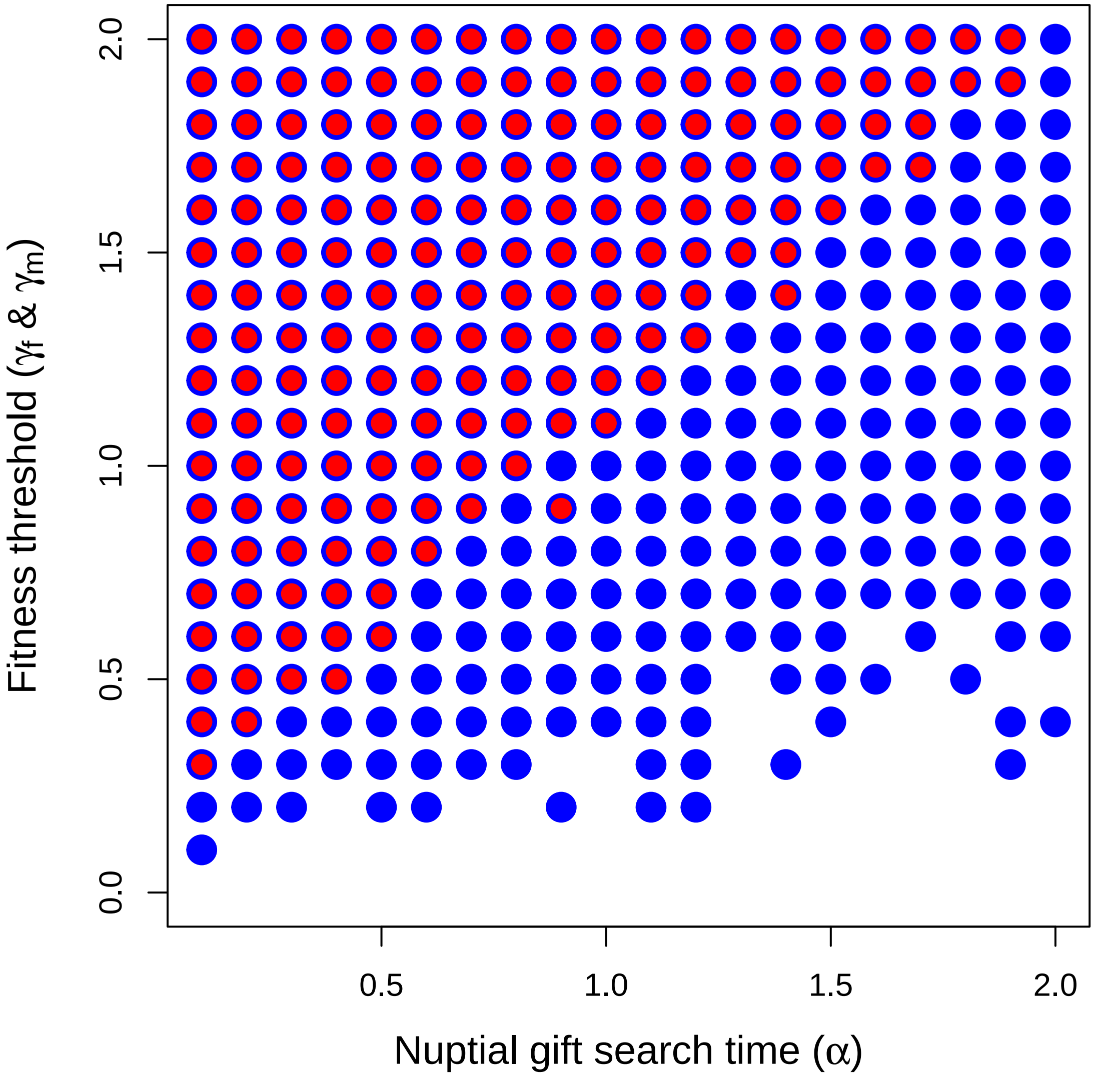


Figure S2.3 ($T_\mathrm{f} = 10.0$): The coevolution of male search and female choosiness as a function of nuptial gift search time ($\alpha$). Points show where the lower 95% confidence interval of female choosiness (red) and male search (blue) exceeds zero, indicating evolution of choosiness or nuptial gift search. Each point includes data from 3000 replicate simulations with identical starting conditions. Up to 3000 interactions occur between individuals in each time step ($\psi = 3$), potentially resulting in a mating interaction. The number of individuals in the population remained at or near carrying capacity of $K = 1000$. Expected female processing time was set to $T_\mathrm{f} = 10.0$ time steps, and $\gamma$ and $\alpha$ values in the range [0.1, 2.0] and [0.0, 2.0], respectively, were used.

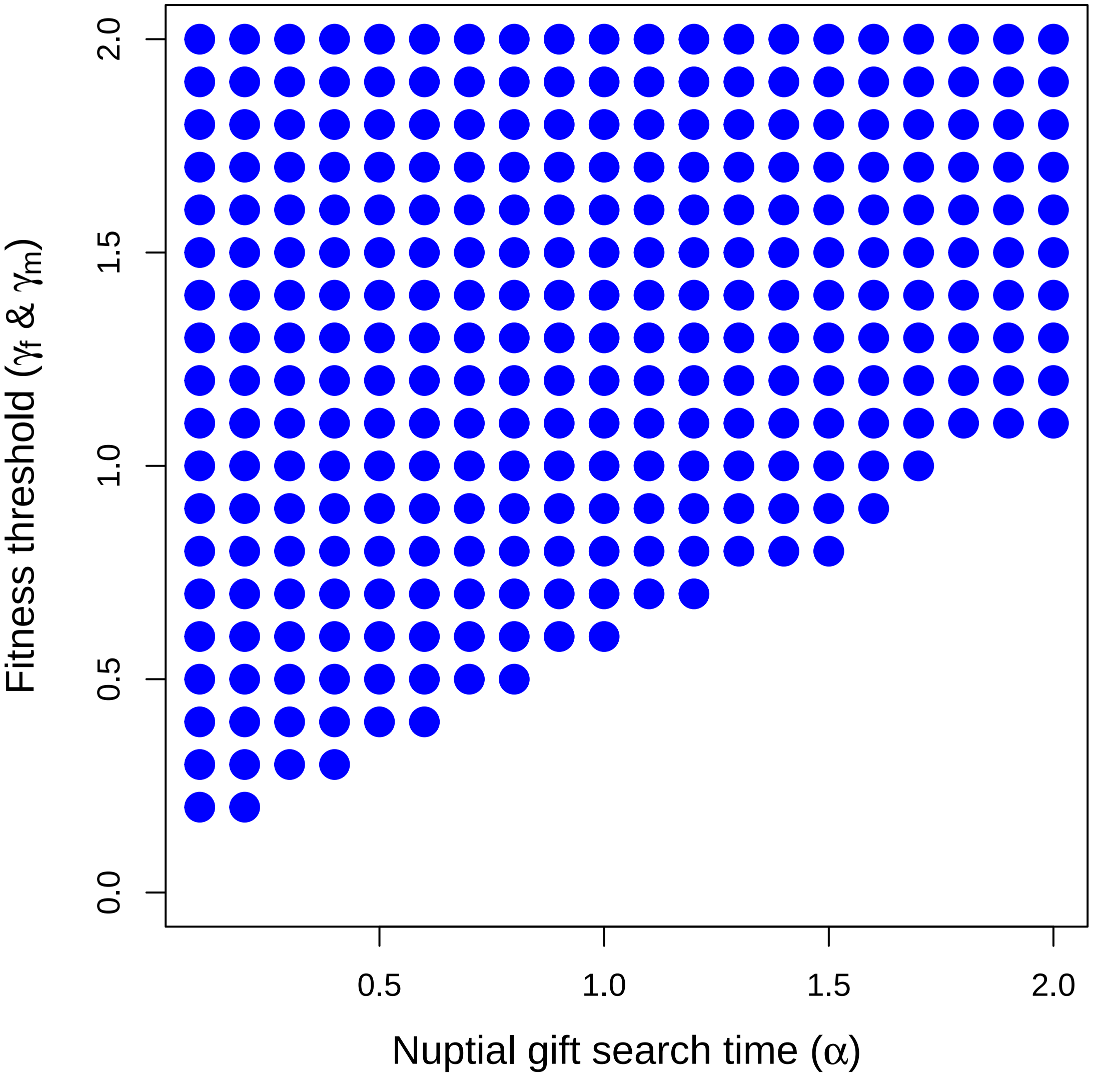


Figure S2.4 ($\psi = 1$): The coevolution of male search and female choosiness as a function of nuptial gift search time ($\alpha$). Points show where the lower 95% confidence interval of where male search (blue) exceeds zero, indicating evolution of choosiness or nuptial gift search. Each point includes data from 3000 replicate simulations with identical starting conditions. Up to 1000 interactions occur between individuals in each time step ($\psi = 1$), potentially resulting in a mating interaction. The number of individuals in the population remained at or near carrying capacity of $K = 1000$. Expected female processing time was set to $T_f = 2$ time steps, and $\gamma$ and $\alpha$ values in the range [0.0, 2.0] and [0.1, 2.0], respectively, were used.

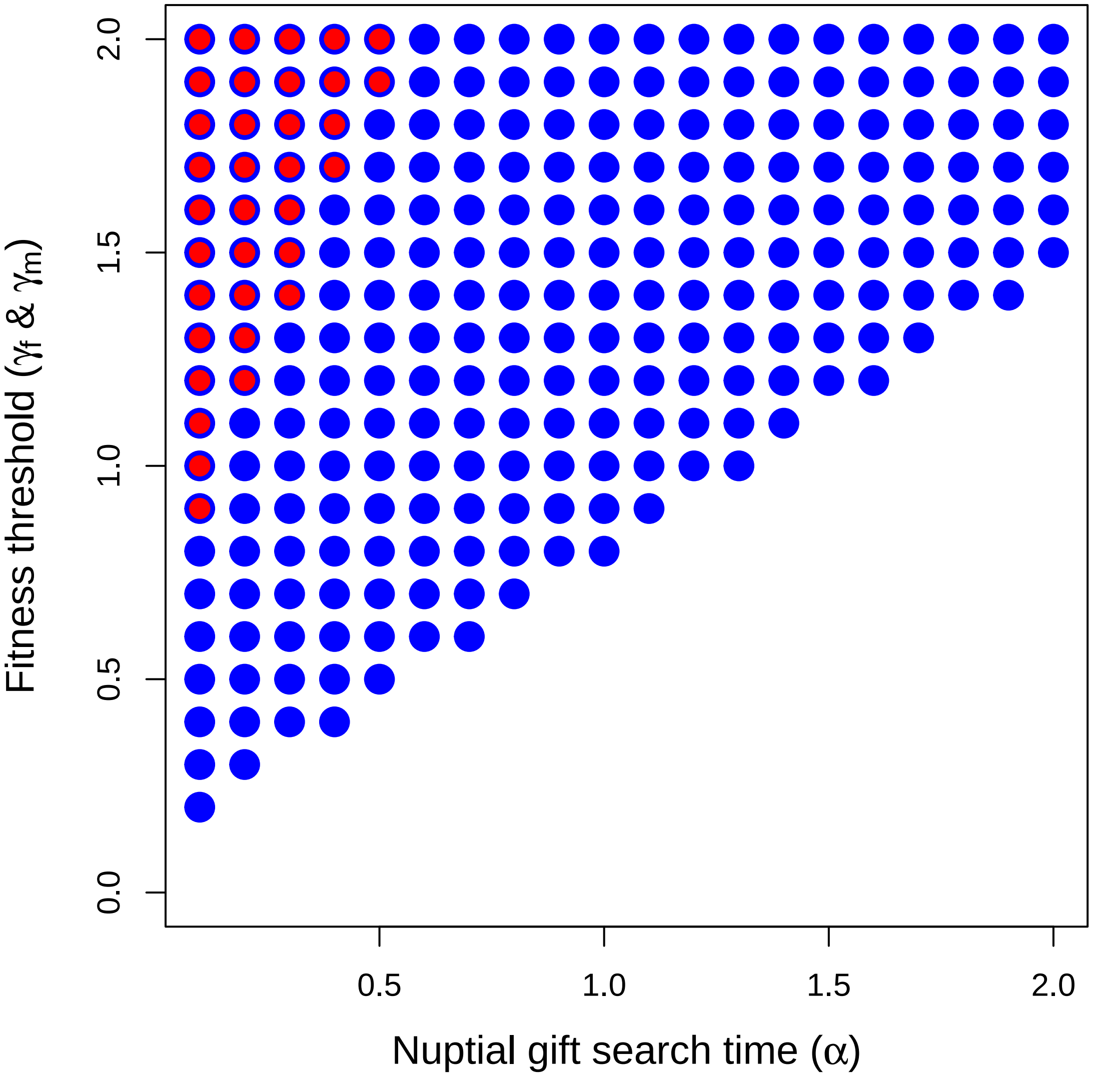


Figure S2.5 ($\psi = 2$): The coevolution of male search and female choosiness as a function of nuptial gift search time ($\alpha$). Points show where the lower 95% confidence interval of where male search (blue) exceeds zero, indicating evolution of choosiness or nuptial gift search. Each point includes data from 3000 replicate simulations with identical starting conditions. Up to 2000 interactions occur between individuals in each time step ($\psi = 2$), potentially resulting in a mating interaction. The number of individuals in the population remained at or near carrying capacity of $K = 1000$. Expected female processing time was set to $T_f = 2$ time steps, and $\gamma$ and $\alpha$ values in the range [0.0, 2.0] and [0.1, 2.0], respectively, were used.

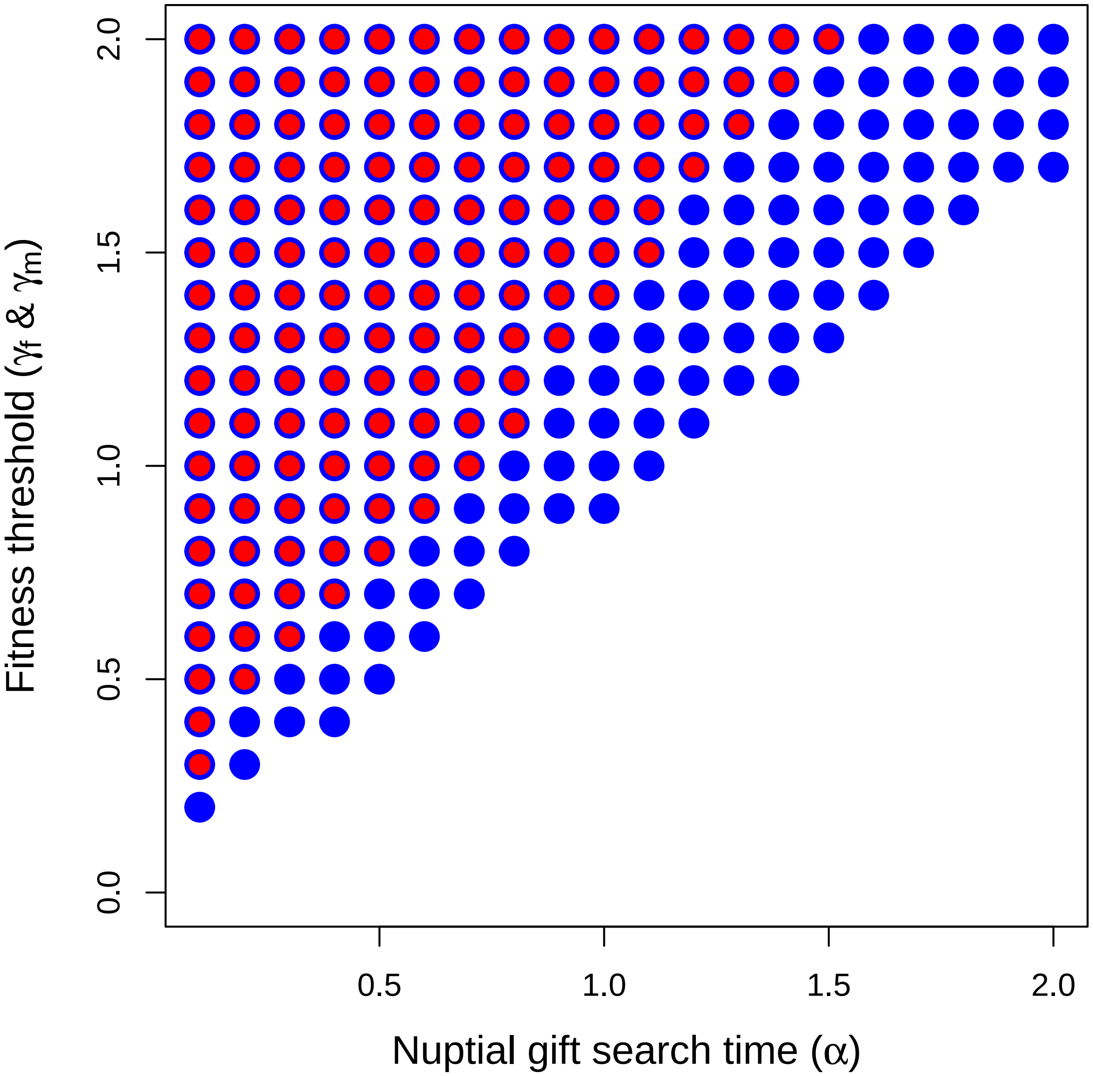


Figure S2.6 ($\psi = 4$): The coevolution of male search and female choosiness as a function of nuptial gift search time ($\alpha$). Points show where the lower 95% confidence interval of where male search (blue) exceeds zero, indicating evolution of choosiness or nuptial gift search. Each point includes data from 3000 replicate simulations with identical starting conditions. Up to 4000 interactions occur between individuals in each time step, potentially resulting in a mating interaction ($\psi = 4$). The number of individuals in the population remained at or near carrying capacity of $K = 1000$. Expected female processing time was set to $T_\mathrm{f} = 2$ time steps, and $\gamma$ and $\alpha$ values in the range [0.0, 2.0] and [0.1, 2.0], respectively, were used.

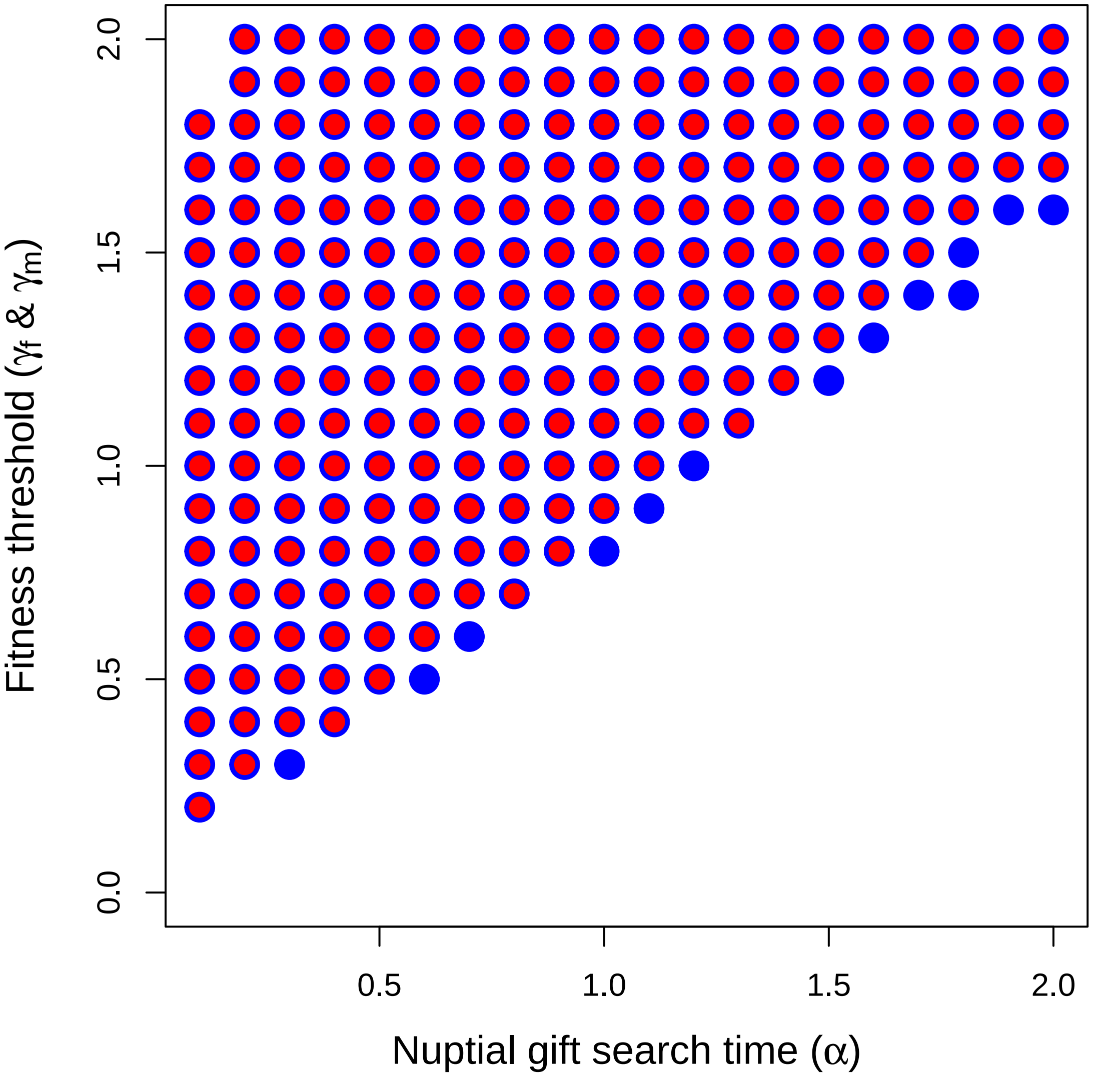


Figure S2.7 ($\psi$ = 6): The coevolution of male search and female choosiness as a function of nuptial gift search time ($\alpha$). Points show where the lower 95% confidence interval of where male search (blue) exceeds zero, indicating evolution of choosiness or nuptial gift search. Each point includes data from 3000 replicate simulations with identical starting conditions. Up to 6000 interactions occur between individuals in each time step, potentially resulting in a mating interaction ($\psi = 6$). The number of individuals in the population remained at or near carrying capacity of $K = 1000$. Expected female processing time was set to $T_f = 2$ time steps, and $\gamma$ and $\alpha$ values in the range [0.0, 2.0] and [0.1, 2.0], respectively, were used.

### S2: Alternative derivation of male fitness threshold

In the main text, we assumed that males made the decision to search or not search for a nuptial gift. The expected length of time for which searching males are expected to remain outside of the mating pool is $E[T_\mathrm{m}] = \alpha$ (see Model). Alternatively, we can assume that males search for a pre-determined period of $T_\mathrm{m}$ and spend this full duration of $T_\mathrm{m}$ in the time-out phase, even if they succeed in finding a nuptial gift (as we assumed in our IBM). The probability that a male obtains a nuptial gift during this time is modelled in Eq. 1,

$$G = 1 - e^{-\frac{1}{\alpha}T_\mathrm{m}}.$$

In Eq. 1, $\alpha$ is the amount of time expected to pass before a male encounters a nuptial gift. We assume that a male will only enter the mating pool with no gift if they are unsuccessful in obtaining a gift, so the probability that a male obtains no gift after $T_\mathrm{m}$ is,

$$L = e^{-\frac{1}{\alpha}T_\mathrm{m}}.$$

We assume that the fitness associated with receiving a nuptial gift versus no nuptial gift are $\lambda(1+\gamma)$ and $\lambda$, respectively. The rate at which males increase their fitness can then be defined as the expected fitness increase from their nuptial gift search divided by $T_\mathrm{m}$ plus the time spent in the mating pool waiting to encounter a mate,

$$W_\mathrm{m} = \lambda \frac{G\left(1+\gamma\right) + L}{T_\mathrm{m} + \left(\frac{\beta+1}{R}\right)}.$$

Our objective now is to determine the conditions under which a focal male increases his fitness by searching for a nuptial gift ($T_\mathrm{m} > 0$) in a population of resident males that do not search ($T_\mathrm{m} = 0$). Females are assumed to exhibit no choosiness for males with versus without nuptial gifts. Under such conditions, male fitness cannot be affected by female choice, so selection to increase $T_\mathrm{m} > 0$ must be based solely on $\alpha$, $\beta$, $R$, and $\gamma$.

To determine under what conditions male inclusive fitness increases with nuptial gift search time, we can differentiate $W_\mathrm{m}$ with respect to $T_\mathrm{m}$,

$$\frac{\partial W_\mathrm{m}}{\partial T_\mathrm{m}} = \lambda \frac{\gamma \left( \frac{\frac{T_\mathrm{m} + \frac{\beta+1}{R}}{\alpha} + 1}{e^{\frac{1}{\alpha}T_\mathrm{m}}} - 1 \right) - 1}{\left(T_\mathrm{m} + \frac{\beta+1}{R}\right)^2}.$$

Because $T_\mathrm{m} = 0$, the above simplifies,

$$\frac{\partial W_\mathrm{m}}{\partial T_\mathrm{m}} = \lambda \frac{\frac{R\gamma(\beta+1)}{\alpha} - R^2}{\left(1+\beta\right)^2}.$$

We can re-arrange the above,

$$\frac{\partial W_\mathrm{m}}{\partial T_\mathrm{m}} = \lambda \frac{R\gamma}{\alpha\left(\beta+1\right)} - \lambda \frac{R^2}{\left(1+\beta\right)^2}.$$

Note that if $R = 0$ or $\lambda = 0$, then, trivially, no change in fitness occurs (since females and males cannot mate or do not produce offspring). Fitness is increased by searching for nuptial gifts when $\gamma$ is high, scaled by the expect search time needed to find a nuptial gift. A second term on the right-hand side is subtracted, which reflects a loss in fitness proportional to the encounter rate of potential mates in the mating pool. The threshold for which male inclusive fitness is not affected by searching for a nuptial gift are found by setting $\partial W_{\mathrm{m}}/\partial T_{\mathrm{m}} = 0$ and solving for $\gamma$ to recover Eq. 2 from the main text,

$$0 = \lambda \frac{R\gamma}{\alpha\left(\beta+1\right)} - \lambda \frac{R^2}{\left(1+\beta\right)^2},$$

$$\lambda \frac{R^2}{\left(1+\beta\right)^2} = \lambda \frac{R\gamma}{\alpha\left(\beta+1\right)},$$

$$\frac{R}{\left(1+\beta\right)^2} = \frac{\gamma}{\alpha\left(\beta+1\right)},$$

$$\frac{R}{\left(1+\beta\right)} = \frac{\gamma}{\alpha},$$

$$\gamma = \alpha \frac{R}{\beta+1}.$$

### S3: Operational sex ratio

We assume that the ratio of males to females is the same upon individual maturation. Consequently, the operational sex ratio $\beta$ will be a function of $R$, $T_{\mathrm{f}}$, and $T_{\mathrm{m}}$ because these parameters determine the density of females and males in the mating pool versus outside of the mating pool. We start with the definition of $\beta$ as being the probability of finding an individual in time-in (Kokko & Monaghan, 2001),

$$\beta = \frac{\int_{t=0}^{\infty} P_{IM}(t)dt}{\int_{t=0}^{\infty} P_{IF}(t)dt}$$

We can substitute the equations for $P_{IM}(t)$ and $P_{IF}(t)$, which define the probabilities of males and females being within the mating pool at time $t$, respectively.

We can therefore calculate $\beta$ as below,

$$\beta = \frac{\left(\frac{\left(\frac{\beta+1}{R}\right)}{T_{\mathrm{m}}+\left(\frac{\beta+1}{R}\right)}\right)}{\left(\frac{\left(R\frac{\beta}{\beta+1}\right)}{T_{\mathrm{f}}+\left(R\frac{\beta}{\beta+1}\right)}\right)}.$$

This can be simplified,

$$\beta = \frac{\left(\beta\left(R+T_f\right)+T_f\right)\left(\beta+1\right)}{\beta\left(R^2T_m+R\right)+\beta^2R}.$$

There is no closed form solution for $\beta$, but a recursive algorithm can be used to calculate $\beta$ to an arbitrary degree of precision.

```
recursive_b <- function(B, D, Tf, Tm, crit = 0.0001, maxit = 9999){
  conv <- 1;
  iter <- 0;
  while(conv > crit & iter < maxit){
    Fe   <- D * (B / (1 + B));
    Me   <- (1 + B) / D;
    Bn   <- (Me / (Tm + Me)) / (Fe / (Tf + Fe));
    iter <- iter + 1;
    conv <- abs(Bn - B);
    B    <- Bn;
  }
  return(list(B = B, conv = conv, iter = iter));
}
```

We used the above function to calculate values of $\beta$ for the analytical model.

### S4: Separate evolution of male search and female choice

We used the individual-based simulation model to unpack the effect of coevolution on the evolution of male search and female choice. Here we replicated the simulations shown in the main text under the condition where only one trait at a time was allowed to evolve and studied how this affected the trait evolution.

First, we submitted a set of simulations wherein male search did not evolve, but was fixed at different values. Next, we ran the same set of simulations wherein male search evolved, but female choice was not possible. The results thus show how each trait evolves in the absence of any coevolution (Fig. S4.1).

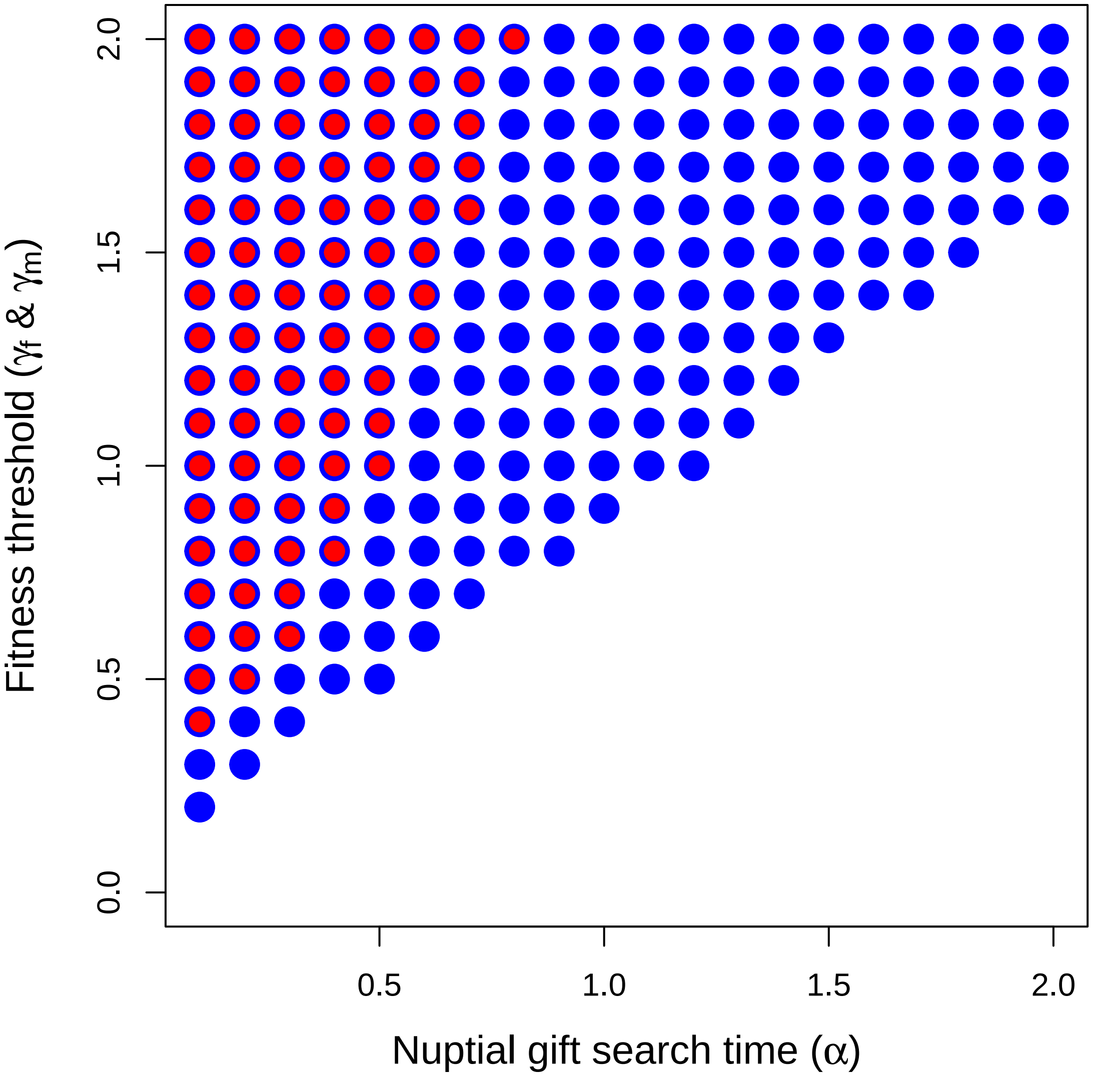


Figure S4.1: The separate evolution of male search and female choosiness as a function of nuptial gift search time. Points show where the lower 95% confidence interval of male search (blue) and female choosiness (red) exceeds zero, indicating evolution of nuptial gift search or choosiness. Each point includes data from $2 \times 3000$ replicate simulations with identical starting conditions. In the first batch, male search was constant and initialized at $T_{\mathrm{m}} = \alpha$, and female choice was evolving. In the second batch, male search was evolving, and there was no option for female choice. The parameters $T_{\mathrm{f}} = 2$, and $\gamma$ and $\alpha$ values were set within the range [0.1, 2.0] and [0.0, 2.0], respectively.

### S5: Male probability of search trait

In the individual-based model of the main text, a focal male $i$ searches for a nuptial gift for a fixed number of Poisson($T_{\mathrm{m}}^{i}$) time steps. The male will remain outside the mating pool for this number of time steps regardless of whether or not he successfully obtains a nuptial gift. Here we instead define male search for nuptial gift as an all-or-nothing strategy to align with our analytical model. Males either search in time-out until they find a nuptial gift, or they do not search at all. In these simulations, $T_{\mathrm{m}}^{i}$ is instead defined as the probability that a focal male $i$ searches for a nuptial gift (in the same way that $\rho^{i}$ is defined as the probability that a focal female $i$ rejects a male without a nuptial gift). For males that decide to search, the time taken until a nuptial gift is obtained is randomly sampled from an exponential distribution with a rate parameter of $\alpha$. This sampling time is then rounded to the nearest integer to determine the number of time steps that a focal male spends outside the mating pool before obtaining a nuptial gift.

Results are qualitatively identical to those in Figure 3 of the main text (Figure S5.1)

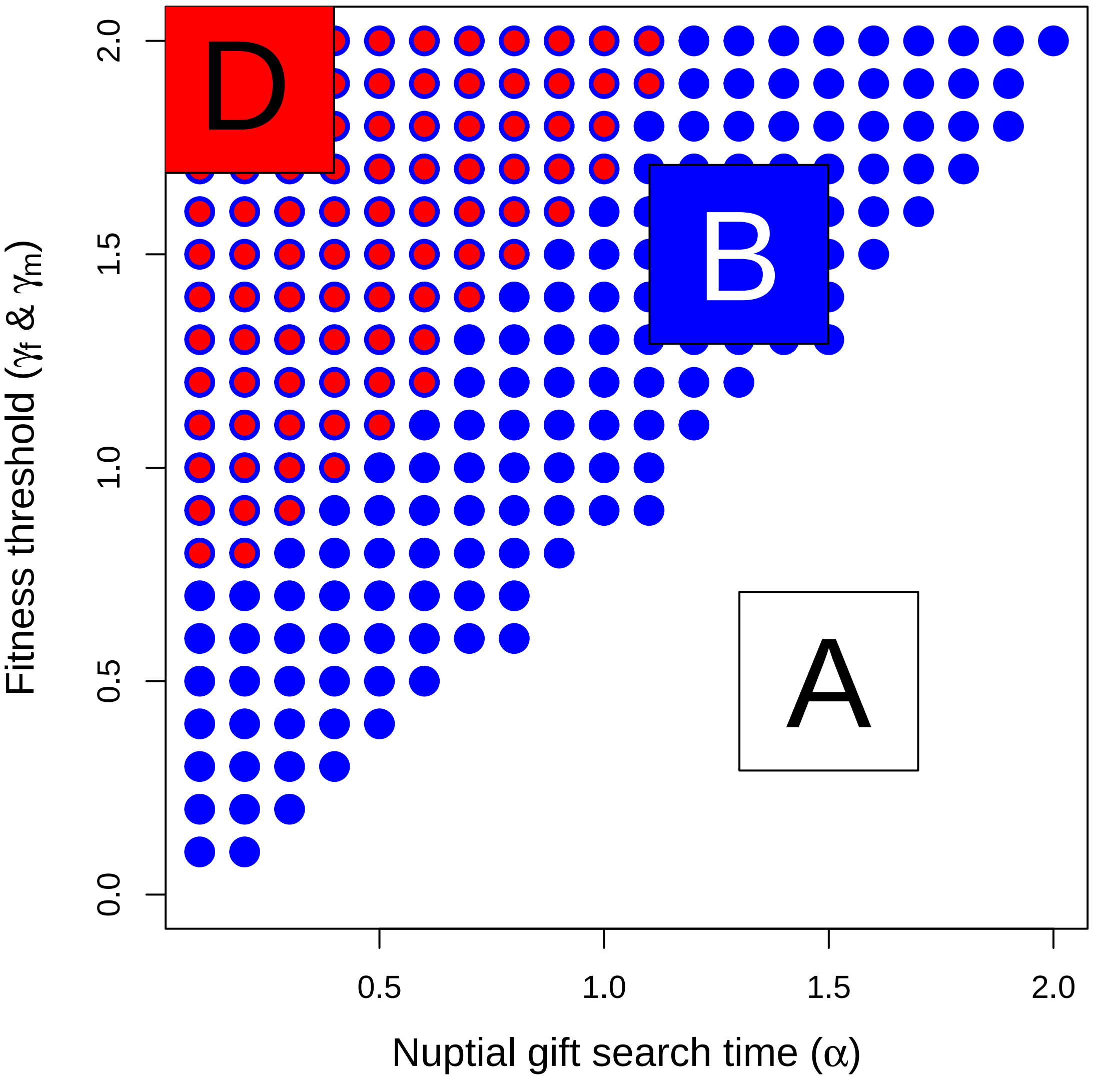


Figure S5.1: Coevolution of male search and female choosiness as a function of nuptial gift search time ($\alpha$) when male strategy is binary: search or do not search. Points show where the lower 95% confidence interval of male search (blue) and female choosiness (red) exceeds zero, indicating evolution of nuptial gift search or choosiness. Each point includes data from 1000 replicate simulations with identical starting conditions. Zones are identified that correspond to areas where males search and females are choosy (D), males search but females are not choosy (B), and males do not search and females are not choosy (A), as also depicted in Figures 2 and 3 in the main text. The number of individuals in the population remained at or near carrying capacity of $K = 1000$. In each time step, up to 3000 total pair-wise interactions occurred. Expected female processing time was set to $T_f = 2$ time steps, and $\gamma$ and $\alpha$ values in the range [0.0, 2.0] and [0.1, 2.0], respectively, were used.
